# Epiclomal: probabilistic clustering of sparse single-cell DNA methylation data

**DOI:** 10.1101/414482

**Authors:** Camila P.E. de Souza, Mirela Andronescu, Tehmina Masud, Farhia Kabeer, Justina Biele, Emma Laks, Daniel Lai, Patricia Ye, Jazmine Brimhall, Beixi Wang, Edmund Su, Tony Hui, Qi Cao, Marcus Wong, Michelle Moksa, Richard A. Moore, Martin Hirst, Samuel Aparicio, Sohrab P. Shah

## Abstract

We present Epiclomal, a probabilistic clustering method arising from a hierarchical mixture model to simultaneously cluster sparse single-cell DNA methylation data and impute missing values. Using synthetic and published single-cell CpG datasets we show that Epiclomal outperforms non-probabilistic methods and is able to handle the inherent missing data feature which dominates single-cell CpG genome sequences. Using a recently published single-cell 5mCpG sequencing method (PBAL), we show that Epiclomal discovers sub-clonal patterns of methylation in aneuploid tumour genomes, thus defining epiclones. We show that epiclones may transcend copy number determined clonal lineages, thus opening this important form of clonal analysis in cancer. Epiclomal is written in R and Python and is available at https://github.com/shahcompbio/Epiclomal.

## Introduction

DNA methylation of the fifth cytosine position (5mC) is a well studied epigenetic mark that has decisive roles in the regulation of cell transcriptional programs [1]. In mammals, 5mC occurs mainly at CpG dinucleotides [2] whose distribution is clustered within regions of the genome called CpG islands (CGIs). Bisulfite mediated conversion of 5mC to uracil, referred to as bisulphite sequencing, has been a key tool for the quantification of genome-wide DNA methylation at single-cytosine resolution. Advances in technology and laboratory protocols have allowed the generation of high-throughput sequencing data of individual cells [3–6]. In particular, single-cell whole-genome bisulfite sequencing (sc-WGBS) techniques have been developed to assess the epigenetic diversity of a population of cells [7, 8]. Because of the limited amount of DNA material, the generated sc-WGBS data are usually sparse, that is, data from a large number of CpG sites are missing and/or are subject to measurement error. Therefore, there is a great need for the development of statistical and computational methods to cluster cells according to their DNA methylation profiles taking into account the large sparsity of the data.

An increasing amount of sc-WGBS data has been generated from various cell types including: mouse embryonic stem cells [9, 10], human hematopoietic stem cells [7, 11], human hepatocellular carcinoma [12], mouse hepatocytes and fibroblasts [13], human and mouse brain cells [14] and human cell lines [15]. In order to assess the epigenetic diversity in these different cell populations, a variety of non-probabilistic methods have been considered. Smallwood et al. [9] propose a sliding window approach to compute methylation rates of CpG sites across the genome followed by complete-linkage hierarchical clustering considering Euclidean distances and the most variable sites. Angermueller et al. [10] compute the mean methylation levels across gene bodies and as in [9] cluster the cells using hierarchical clustering and only the most variable genes. Farlik et al. [11] cluster cells based on the average methylation over different sets of transcription factor binding sites using also hierarchical clustering. Gravina et al. [13] consider the sliding window approach of [9] to compute methylation rates and use principal component analysis to visually assess clusters of cells. Hou et al. [12] consider CpG-based Pearson correlation between pairs of cells followed by hierarchical clustering. Luo et al. [14] first apply a hierarchical clustering method called BackSPIN [16] to bin-based methylation rates followed by cluster merging using gene body methylation levels. Mulqueen et al. [15] use NMF (non-negative matrix factorization, [17]) and t-SNE [18] for dimensionality reduction, followed by DBSCAN [19] for clustering. Hui et al. [7] proposed PDclust, a genome-wide pairwise dissimilarity clustering strategy that leverages the methylation states of individual CpGs. Recently, Kapourani and Sanguinetti [20] proposed a probabilistic clustering method based on a hierarchical mixture of probit regression models and focused their evaluation on missing CpG data imputation. Angermuller et al. [21] also proposed a deep learning approach for CpG missing data imputation, but which does not address the clustering problem.

Despite the considerable diversity in clustering approaches, there is still a great need for probabilistic, model-based approaches to simultaneously cluster sc-WGBS data while also inferring the missing methylation states. Because such methods allow for statistical strength to be borrowed across cells and neighbouring CpGs, we surmise they should result in more robust inference than non-probabilistic methods. Moreover, no comparative studies to rigorously evaluate the performance of different methods on a wide variety of data sets has been undertaken.

In this work, we propose Epiclomal, a probabilistic algorithm to cluster sparse CpG-based DNA methylation data from sc-WGBS. Our approach is based on a hierarchical mixture model (see graphical models in Figure 1), which pools information from observed data across all cells and neighbouring CpGs to infer the cell-specific cluster assignments and their corresponding hidden methylation profiles. Epiclomal is part of a novel comprehensive statistical and computational framework (Figure 2) that includes data pre-processing, different clustering methods corresponding to previously proposed approaches [7, 9, 10, 12, 15], plotting and quantitative performance evaluation measures to analyze the results. We use our framework to present an assessment of clustering methods over previously published and synthetic data sets, plus a novel large scale sc-WGBS data set from breast cancer xenografts [22, 23] generated using state of the art methodology [7].

**Fig 1.**
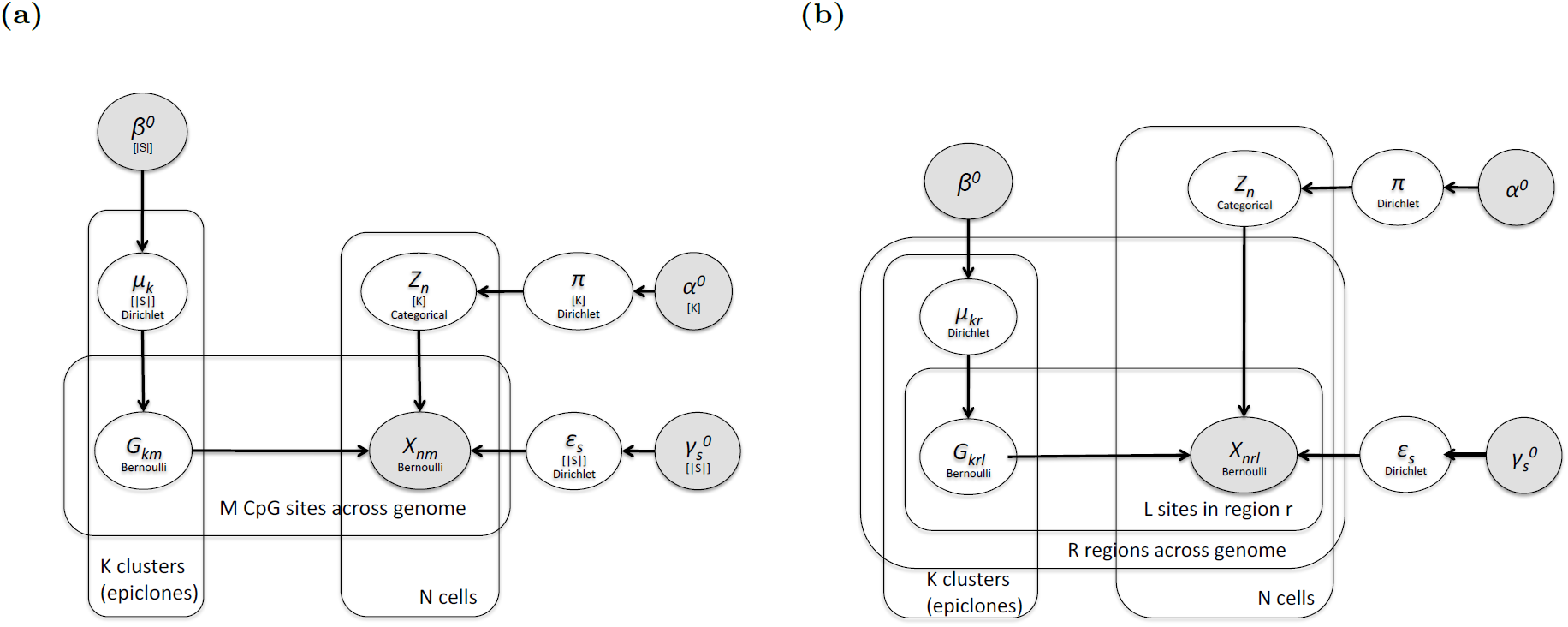
(a) EpiclomalBasic and (b) EpiclomalRegion graphical models. In (a) the shaded node *X*_*nm*_ denotes the observed methylation state at CpG site *m* of cell *n*. In (b) we take into account the region location of each CpG and let the shaded node *X*_*nrl*_ denote the observed methylation state at CpG site *l* of region *r* of cell *n*. Both *X*_*nm*_ and *X*_*nrl*_ take value in 𝒮 = {unmethylated, methylated} or simply 𝒮 = {0, 1}. In (a) and (b), the unshaded *Z*_*n*_ node corresponds to the latent variable (taking value in {1, …, *K*}) indicating the true cluster population (epiclone) for cell *n*. The *G*_*km*_ and *G*_*krl*_ unshaded nodes in (a) and (b), respectively, are the latent variables taking value in 𝒮 that correspond to the true hidden CpG epigenotypes for each epiclone *k*. The unshaded *µ, π* and *ϵ* nodes in both (a) and (b) correspond to the unknown model parameters, which under the Bayesian paradigm have prior distributions with fixed hyperparameters described in the shaded nodes with the 0 superscript. The distribution assumed for each variable/parameter is written within its node. The edges of the graphs depict dependencies. The plates depict repetitions. In EpiclomalBasic (a), true hidden epigenotypes share the same probability distribution across all CpG sites in the same epiclone (*G*_*km*_ ∼ Bernoulli(*µ*_*k*_)). In EpiclomalRegion (b), true hidden epigenotypes follow a Bernoulli distribution with probability parameters that vary across regions (*G*_*krl*_ ∼ Bernoulli(*µ*_*kr*_)).

**Fig 2.**
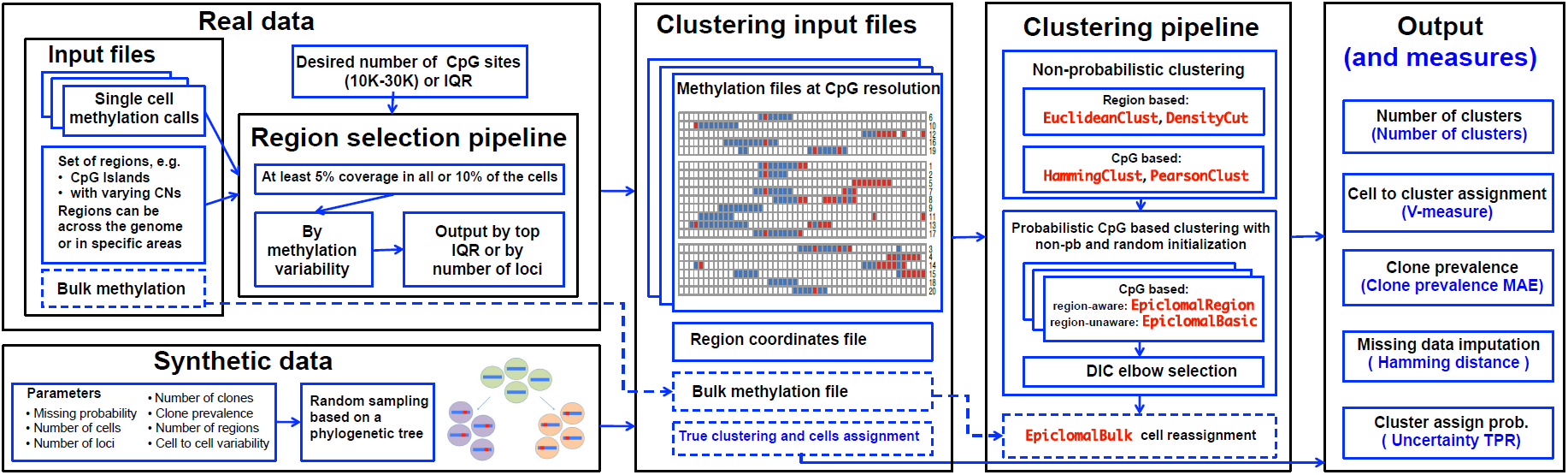
The three components of our proposed framework. *Input data and pre-processing:* data from regions of interest are extracted from methylation call files, which can be filtered keeping only data from regions with a desired amount of missing data and methylation level IQR. A synthetic data pipeline is also provided to simulate data under different parameters. *Clustering:* cells are clustered using different non-probabilistic clustering methods, whose results will be then used as initial values for Epiclomal methods. *Output and performance measures:* different metrics are provided to evaluate the output of each method when true cluster assignments are known.

## Results

### Overview of Epiclomal

Epiclomal is a clustering method based on a hierarchical mixture of Bernoulli distributions. We are given a sparse matrix of *N* rows (cells) and *M* columns (CpG sites), in which each entry is either 0 (unmethylated), 1 (methylated) or missing. The distribution of the observed data *X*_*nm*_ for each CpG site *m* from cell *n* depends on the latent cell-specific cluster assignment *Z*_*n*_ and corresponding true hidden methylation state (epigenotype) at that CpG, *G*_*km*_ (Figure 1a). We use a Variational Bayes (VB) algorithm (Supplementary Material Section 2) with random and informed initializations to infer not only cell-to-cluster assignments but also the true hidden cluster-specific epigenotypes *G*_*k*1_, …, *G*_*kM*_ for each cluster *k*, for *k* = 1, …, *K*. We run Epiclomal considering *K* from 1 to a maximum number of possible clusters and choose the best *K* along with the best clustering assignments as the combination that minimizes the deviance information criterion (DIC, [24]) via an elbow plot type of selection procedure (Supplementary Figure 1).

Epiclomal has two variants: EpiclomalBasic (Figure 1a) and EpiclomalRegion (Figure 1b). While EpiclomalBasic imposes less structure on the model by assuming the true hidden methylation states share the same distribution across all the CpG sites considered, EpiclomalRegion allows their distribution to vary across genomic functional regions such as CGIs to better reflect expected behaviour in the real data. Bulk data can be used to reassign cells to the EpiclomalRegion clusters using an algorithm that stochastically reassigns cells to clusters while trying to best match the cumulative CpG states of all cells to the corresponding bulk CpG state. This extension is called EpiclomalBulk (Section 2.3 of the Supplementary Material).

Epiclomal is then incorporated into the computational framework presented in Figure 2 and described in what follows.

### Overview of proposed framework

#### Input data and pre-processing

Our framework (Figure 2) can take as input either real or synthetic data. For real data we take files with CpG methylation calls across the genome from individual cells and extract data from defined regions of interest (e.g., CGIs, gene bodies, differentially methylated regions, etc). Since not all CpG sites exhibit variation and, therefore, are uninformative for clustering, our framework optionally allows selection of specific regions. One can then consider the data from all regions of interest or apply our region selection pipeline to use data from a subset of those regions. Our proposed selection pipeline first keeps the regions with at least 5% coverage in all or 10% of cells and then selects regions with the most variable methylation levels across cells (via interquartile range - IQR), optionally controlling for a desired number of CpG loci. If available, our framework also takes as input bulk methylation data that can be used to inform inference.

For synthetic data we provide a pipeline that generates single-cell methylation data considering various parameters (e.g., missing proportion, number of cells, number of loci, etc.), assuming true cluster methylation profiles arise from a phylogenetic process with loci changing methylation states at each new cluster generation (Supplementary Material Section 3). Synthesis is motivated by tumour clonal composition theory [23, 25] where clonal sub-populations arise from a hierarchical ancestor-descendant phylogenetic process. We note this difference from the independence assumption among epiclones in the EpiClomal model.

#### Cluster initialisation

Given methylation calls and genomic coordinates of retained regions, we first cluster cells considering different non-probabilistic methods, whose results will be then used as initial values for Epiclomal as well as for comparison purposes. We deploy two types of non-probabilistic clustering methods: region and CpG based (Supplementary Methods Section 1). In the region-based approaches we cluster cells considering as input the mean methylation level of each region: EuclideanClust uses hierarchical clustering with Euclidean distances; and DensityCut [26] is a density-based clustering method applied after dimensionality reduction using principle component analysis. In the CpG-based approaches we consider the methylation state of each individual CpG: HammingClust and PearsonClust use hierarchical clustering with Hamming distances and Pearson correlation values, respectively. To find the optimal number of clusters, DensityCut includes its own automatic method, whereas the hierarchical clustering methods use the Calinski-Harabasz (CH) index [27].

Our pipeline runs Epiclomal using the results of the non-probabilistic methods as initial values along with a set of random initials, and chooses the best configuration as explained in the “Overview of Epiclomal” section.

#### Output and performance measures

For all clustering methods we output predictions of cell-to-cluster assignments, number of clusters, and cluster (or epiclone) prevalences (i.e., the proportion of cells assigned to each cluster). In addition, for Epiclomal we obtain the estimated missing CpG values and the cell-to-cluster assignment posterior probabilities.

When ground-truth clustering is available, we also output a performance evaluation measure for each of the five aforementioned predictions (Figure 2 and Supplementary Material Section 4). V-measure evaluates the cell-to-cluster assignments [28] and it is a score between zero and one, where one stands for perfect clustering and zero for random cell-to-cluster assignments. The V-measure captures the homogeneity and completeness of a clustering result. To satisfy the homogeneity criterion, a clustering procedure must assign only those cells that are members of a single group to a single cluster. Completeness is satisfied if all of those cells that are members of a single group are assigned to a single cluster. The harmonic mean of homogeneity and completeness gives rise to the V-measure, and even a small percentage of misclassified cells can significantly affect it. We also report the predicted number of clusters and the mean absolute error (MAE) between true and predicted cluster prevalence values. In addition, when applying Epiclomal on synthetic data, we consider the Hamming distance as the proportion of discordant entries between true and inferred vectors of methylation states. We also compute for Epiclomal the uncertainty true positive rate of cluster assignment probabilities, that is, how well the uncertainty is estimated for cells whose membership is unclear due to missing data.

### Epiclomal outperforms other methods on synthetic data

To evaluate the performance of our proposed methods over a wide range of characteristics, we generated a large number of synthetic datasets and applied our Epiclomal approaches (EpiclomalRegion, EpiclomalBasic and EpiclomalBulk), as well as the four non-probabilistic methods (EuclideanClust, DensityCut, HammingClust and PearsonClust) to each generated data set.

We considered several experiments, where in each one we varied one of seven parameters while keeping the others fixed as indicated in Table 1. For each setting, we generated 30 input datasets and ran Epiclomal with a total of 300 informed and random VB initialisations. Then, we computed the V-measure along with the other quantities included in our framework to assess method performance.

**Table 1.** The varying parameters and their ranges for the synthetic data simulation. For each experiment, we varied one parameter and kept the other ones fixed. Note that varying the number of regions is equivalent to varying region size as the total number of loci is fixed. Unless otherwise specified, the fixed parameters are: missing proportion 0.8, region size 100, number of cells 100, proportion of cell to cell variability 0, number of epiclones 3, equal epiclone prevalences (1/3) and number of loci 10 000. For the cell-to-cell variability experiment (Figure 4c) we used 25 regions in order to have a larger number of loci that differ between clusters. For the number of epiclones experiment (Figure 4d) and the epiclone prevalence experiment (Figure 4e) we used 500 cells in order to allow for enough cells to be represented in each case. For the number of loci experiment (Figure 4f) we also varied the number of regions in order to keep differences among clusters fixed (e.g., 50 regions for 5 000 loci, 5 000 regions for 500 000 loci)

| Varying parameter | Varying range |
| --- | --- |
| Missing proportion | 0.5 to 0.95 |
| Number of regions | 25 to 200 |
| Number of cells | 12 to 2500 |
| Cell-to-cell variability | 0 to 0.3 |
| Number of clusters (epiclones) | 1 to 10 |
| epiclone prevalence | balanced to very unbalanced |
| Number of loci | 5 000 to 500 000 |

Figure 3 shows the results when we vary the proportion of data missing from 0.5 to 0.95. Our proposed probabilistic Epiclomal methods give better or comparable V-measures (panel a) with overall more correct number of clusters (*K* = 3, panel b) than the non-probabilistic methods, which tend to overestimate (EuclideanClust) or underestimate (PearsonClust, HammingClust and DensityCut) the number of clusters. PearsonClust and HammingClust fail to produce results in the case of 0.95 missing proportion. Using bulk data via EpiclomalBulk shows improvement in estimating cluster prevalences, especially when the missing data proportion is large (0.9 and 0.95, panel c). The cluster assignment uncertainty is well estimated by EpiclomalRegion for up to 0.7 missing proportion; however, it drops rapidly for 0.8 and 0.9 missing proportion (panel d). EpiclomalRegion better infers the vectors of cluster-specific methylation profiles than EpiclomalBasic with even smaller hamming distances obtained after adjusting the imputed states for CpGs with no observed data across an entire cluster (see Supplementary Figure 2 and Supplementary Material Section 2.4).

**Fig 3.**
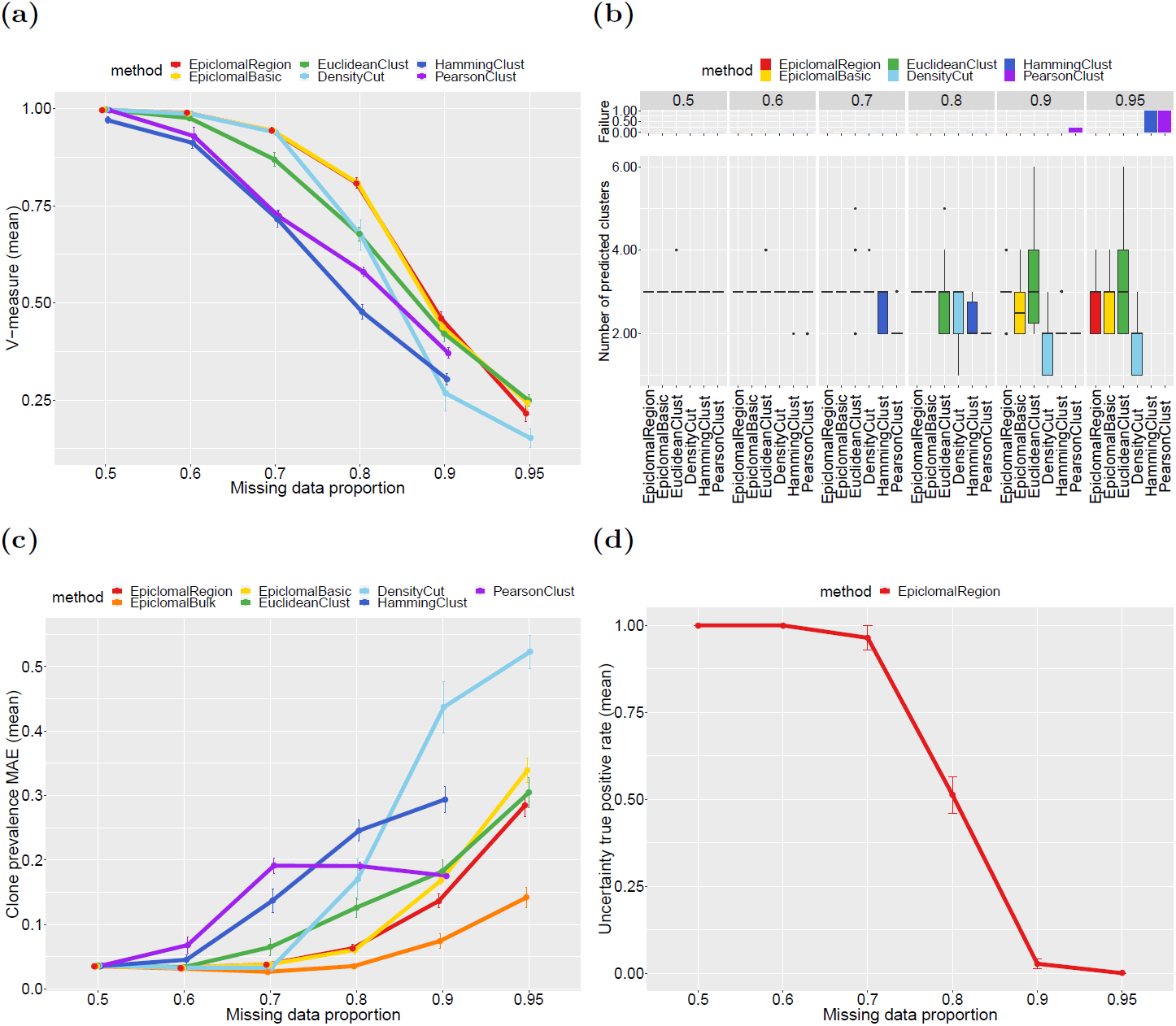
Simulation results when varying the missing data proportion. We report mean results produced by Epiclomal and the non-probabilistic methods taken over 30 randomly generated synthetic datasets: (a) V-measure; (b) Number of predicted clusters (true is 3), the top panel shows the proportion of data sets for which a method failed to produce a result; (c) Epiclone prevalence MAE (mean absolute error); (d) Uncertainty true positive rate. The vertical bars correspond to one standard deviation above and below the mean value. Source data for this figure are provided in file SourceDataFig3.xlsx

Figure 4 shows that Epiclomal results in better V-measure when compared with the non-probabilistic methods in all the remaining experiment scenarios with fixed missing proportion of 0.8 (see also Supplementary Figures 3 to 8). All methods perform worse when the problem is more difficult, such as when increasing the number of regions (therefore decreasing the number of different loci among clusters - Figure 4 panel a) or increasing the cell-to-cell variability (panel c). Increasing the number of cells (panel b) does not improve the V-measure values, except for DensityCut, but it does reduce their variability. The Epiclomal methods are more robust to the increase in the number of epiclones (panel d) and change in epiclone prevalences (panel e). When increasing the number of loci (panel f) the performance of HammingClust and PearsonClust remains somewhat constant, while the other methods show a decreasing pattern in performance. However, the Epiclomal methods still perform better for all number of loci considered than all the other methods. Therefore, this provides support to a strategy for selecting a smaller number of loci (under 50 000) in order to keep the true signal and eliminate noise when analyzing a real data set.

**Fig 4.**
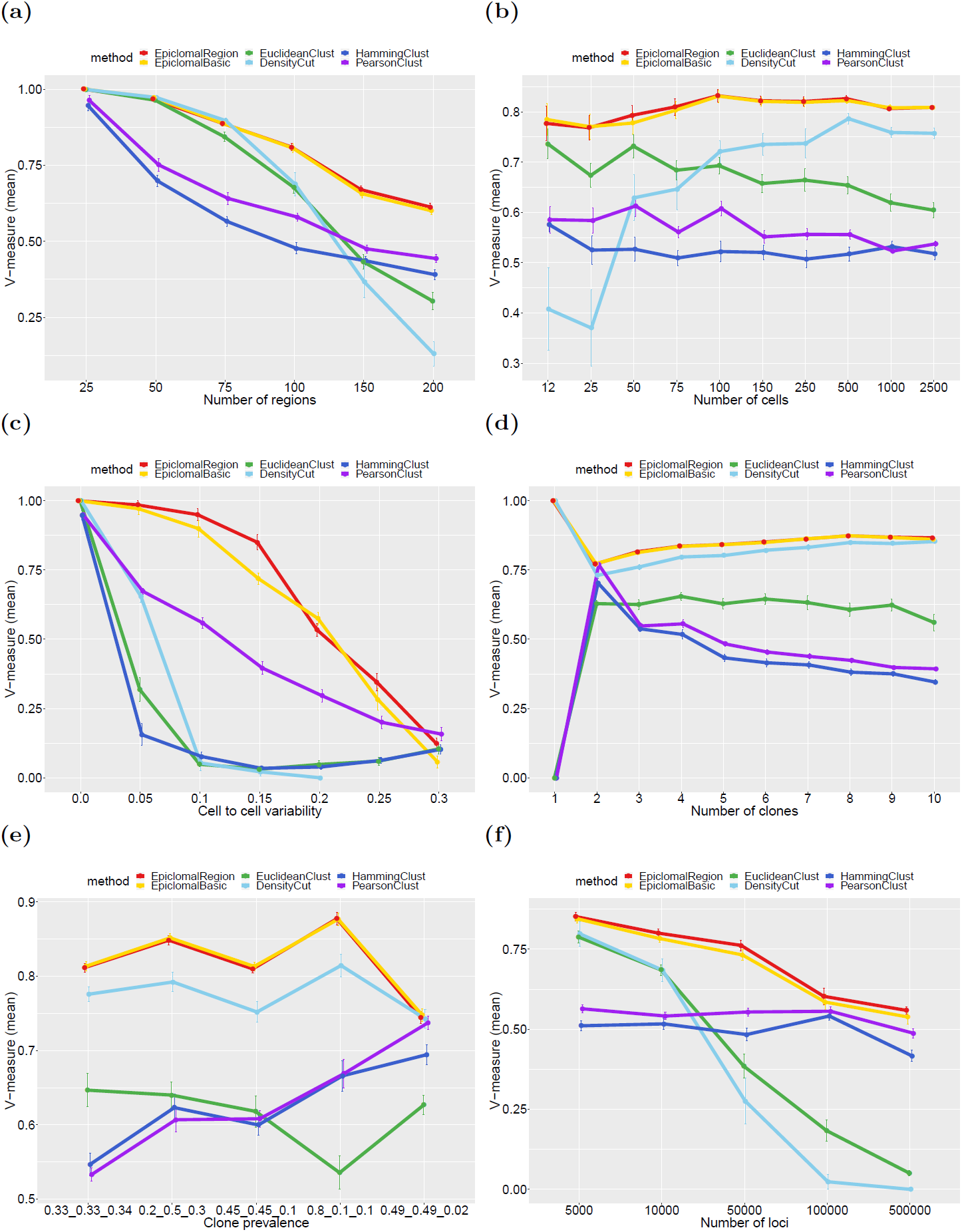
Additional simulation results. We report mean V-measures produced by Epiclomal and the non-probabilistic methods taken over 30 randomly generated synthetic data sets, when we vary: (a) the number of regions, (b) the number of cells, (c) the cell-to-cell variability, (d) the number of clones, (e) the cluster prevalence, (f) the number of loci. The vertical bars correspond to one standard deviation above and below the mean value. The Epiclomal methods outperform the other methods in all cases. Source data for this figure are provided in file SourceDataFig4.xlsx

### Epiclomal recapitulates methylation subgroups from public datasets

We further assessed the performance of our methods on three published sc-WGBS datasets [9, 11, 12] and compared with the clustering results reported in each paper. Experimental validation of epiclones is often difficult, therefore when working with cells from different known types or treatment conditions authors expected their clusters to somewhat reflect the epigenetic diversity of those types [9, 11]. In [12], there were no predefined cell subpopulations, however the authors consider gene expression and copy number changes to further support their findings. Table 2 shows that these datasets display a variety of characteristics, with missing data proportions varying from 0.54 to 0.98.

**Table 2.** A summary of the real data sets used in this work. Column descriptions (in order of appearance) are as follows: (1) data set names corresponding to four published data sets and one new in-house data set; (2) type of cells in each data set; (3) number of cells considered for each data set, these vary from tens to thousands of cells; (4) number of clusters, as reported in the respective published papers or as expected for the InHouse data set; (5) genomic functional regions considered for each data set, these were the same as in the original papers when applicable, TFBS stands for Transcription Factor Binding Sites; (6) missing proportion for each data set considering the 10 000 loci filtered input, varying from 0.69 to 0.94; (7) number of loci for the largest input data sets obtained by including all regions with methylation IQR ≥ 0.01, these vary from 1/4 million to 1 million CpG sites; (8) missing proportion for the largest input datasets, these vary from 0.54 to 0.98.

| Data set | Cell type | # cells | # clusters | Regions | Miss 10K | Nloci IQR $\geq .01$ | Miss IQR $\geq .01$ |
| --- | --- | --- | --- | --- | --- | --- | --- |
| Smallwood2014 [9] | mouse embryonic stem cells | 32 | 2 | CpG Islands | 0.69 | 786 620 | 0.54 |
| Hou2016 [12] | human hepatocellular carcinomas | 25 | 2 | CpG Islands | 0.87 | 255 136 | 0.90 |
| Farlik2016 [11] | human hematopoietic cells | 122 | 6 | TFBS | 0.89 | 512 153 | 0.98 |
| InHouse (3 patients) | human xenografted cancer cells | 558 | 3 | CpG Islands | 0.82 | 1 019 956 | 0.79 |

In what follows we present the results (Figure 5 and Supplementary Figure 9) of applying our framework (Figure 2) to these published datasets considering a filtered input of about 10 000 loci (see Methods for details on pre-processing real data). Supplementary Figure 9 also shows the results for larger input datasets. The Smallwood2014 dataset [9] comprises 32 mouse embryonic stem cells, where 20 cells were cultured in a regular serum medium and 12 cells in a 2i medium inducing hypomethylation. Figure 5a shows a very good agreement between the clusters inferred by EpiclomalRegion and the ones obtained by [9] with only one discordant cell (V-measure 0.82, Figure 5e and Supplementary Figure 10). PearsonClust correctly clustered all cells (V-measure = 1, Figures 5e), while the other non-probabilistic methods misclassified one or two cells (Figure 5e).

**Fig 5.**
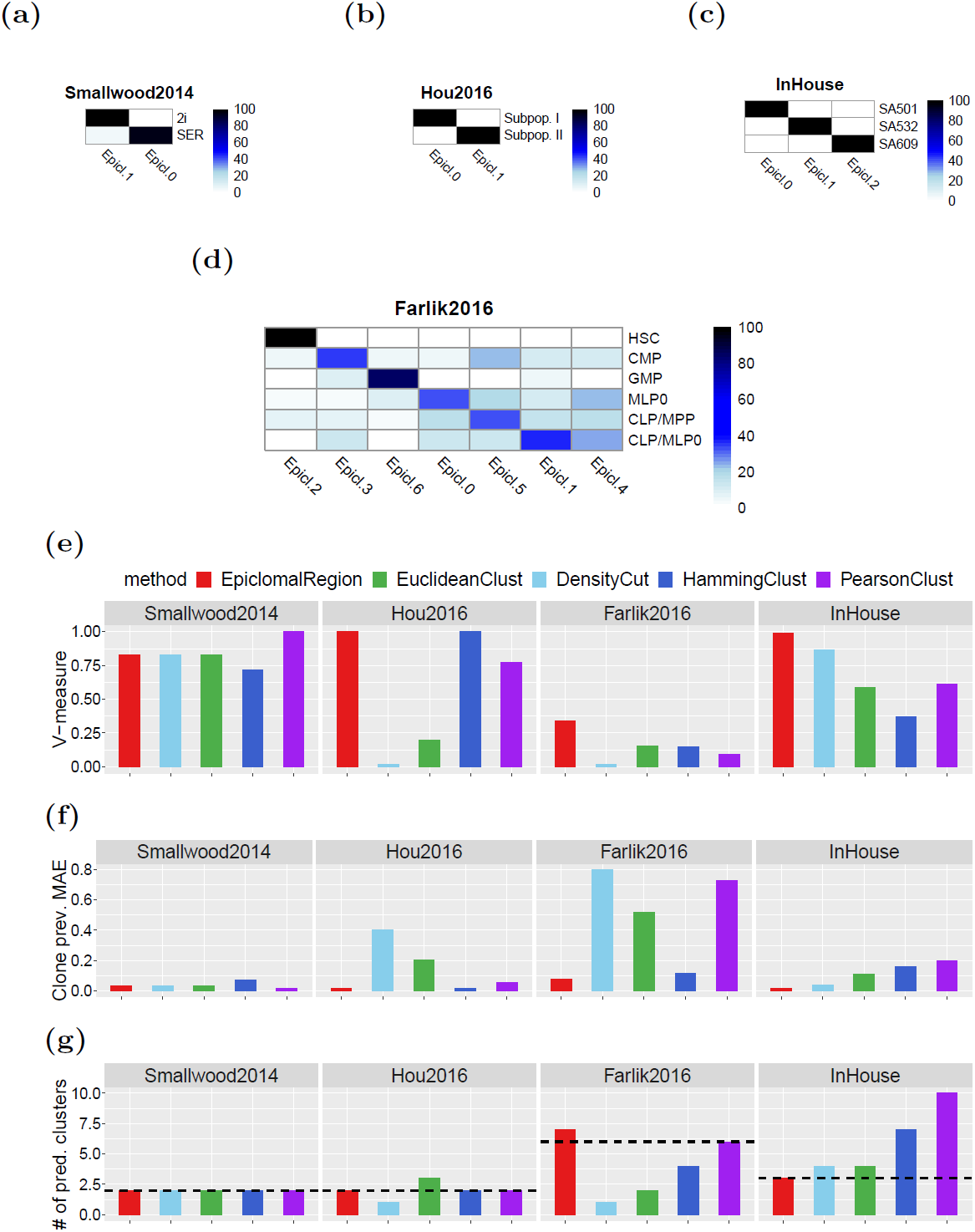
Results for real data sets. (a)-(d) Co-clustering between the real data clusters on the rows and EpiclomalRegion predictions on the columns. Each entry *a*_*ij*_ is the percentage of cells in true class *i* that are present in predicted cluster *j*, rows sum up to 100%. A perfect agreement would result in a square matrix with black diagonal. Rows are ordered as in the original papers; columns are ordered to best match the corresponding true class. (e) V-measures comparing the cell assignments with the true assignments, the higher the better. (f) Clone prevalence mean absolute error, comparing the proportions of clones with the true proportions, the lower the better. (g) Number of predicted clusters (filter using 10 000 loci). The horizontal dashed lines correspond to the true number of clusters; the closer to this line the better. Source data for this figure are provided in file SourceDataFig5.xlsx

The Hou2016 dataset [12] contains 25 cells from a human hepatocellular carcinoma tissue sample. We compare our results with the two subpopulations identified by [12] based not only on DNA methylation but also on copy number and gene expression data. EpiclomalRegion correctly assigned all cells to their corresponding subpopulation (V-measure = 1, Figures 5b, 5e and Supplementary Figure 11).

The Farlik2016 data set [11] contains different types of human hematopoietic cells, totalizing 122 cells. We compared our results with the six clusters found by Farlik et al. [11], comprising hematopoietic stem cells (HSC) and progenitor cell types (myeloid, multipotent and lymphoid progenitor cells). EpiclomalRegion resulted in a V-measure of 0.34, with 7 predicted clusters (Figures 5d, e and g). As stated before, the V-measure can be significantly affected by a small percentage of misclassified cells, therefore, even though the V-measure is low, we observe in Figure 5d a good agreement between Epiclomal clustering and the clustering reported by Farlik et al.

Figures 5e-g show that EpiclomalRegion generally outperforms the non-probabilistic methods on V-measure, clone prevalence mean absolute error and number of correctly predicted clusters.

### Epiclomal reveals copy number dependent and copy number independent epiclones in breast cancer

Having verified the performance of Epiclomal on synthetic data and public domain datasets, we set out to perform epiclone group discovery on single cell epigenomes generated in house on a range of patient-derived breast tumour xenografts. First, to illustrate scalability of Epiclomal with aneuploid single cell cancer epigenomes, we analysed 558 tumour xenograft single epigenomes from two patients with triple-negative breast cancer and one patient with ER+PR-Her2+ breast cancer (Supplementary Table 1), sequenced using the PBAL method [7]. We considered as ground-truth three major clusters, one per patient, for the following two reasons: (1) clustering by cancer type (two clusters) would be incorrect due to major observed epigenotype differences between the two triple-negative breast cancer patients SA501 and SA609, (Figure 6), and (2) inter-patient CpG methylation differences are expected to dominate over minor intra-patient CpG methylation variations [29]. EpiclomalRegion correctly classified all cells except one very sparse cell (V-measure = 0.99, Figure 5c). Methylation differences can be observed between the three patients (Figure 6), with SA609 having a very different methylation profile than the other two. Note that two different experimental plates of very different missing proportions for SA532 (panel a) resulted in visually distinct subclusters in panel b, potentially affecting density-based approaches. Indeed, DensityCut clustered these plates into two different clusters, yet Epiclomal was robust to this batch effect.

**Fig 6.**
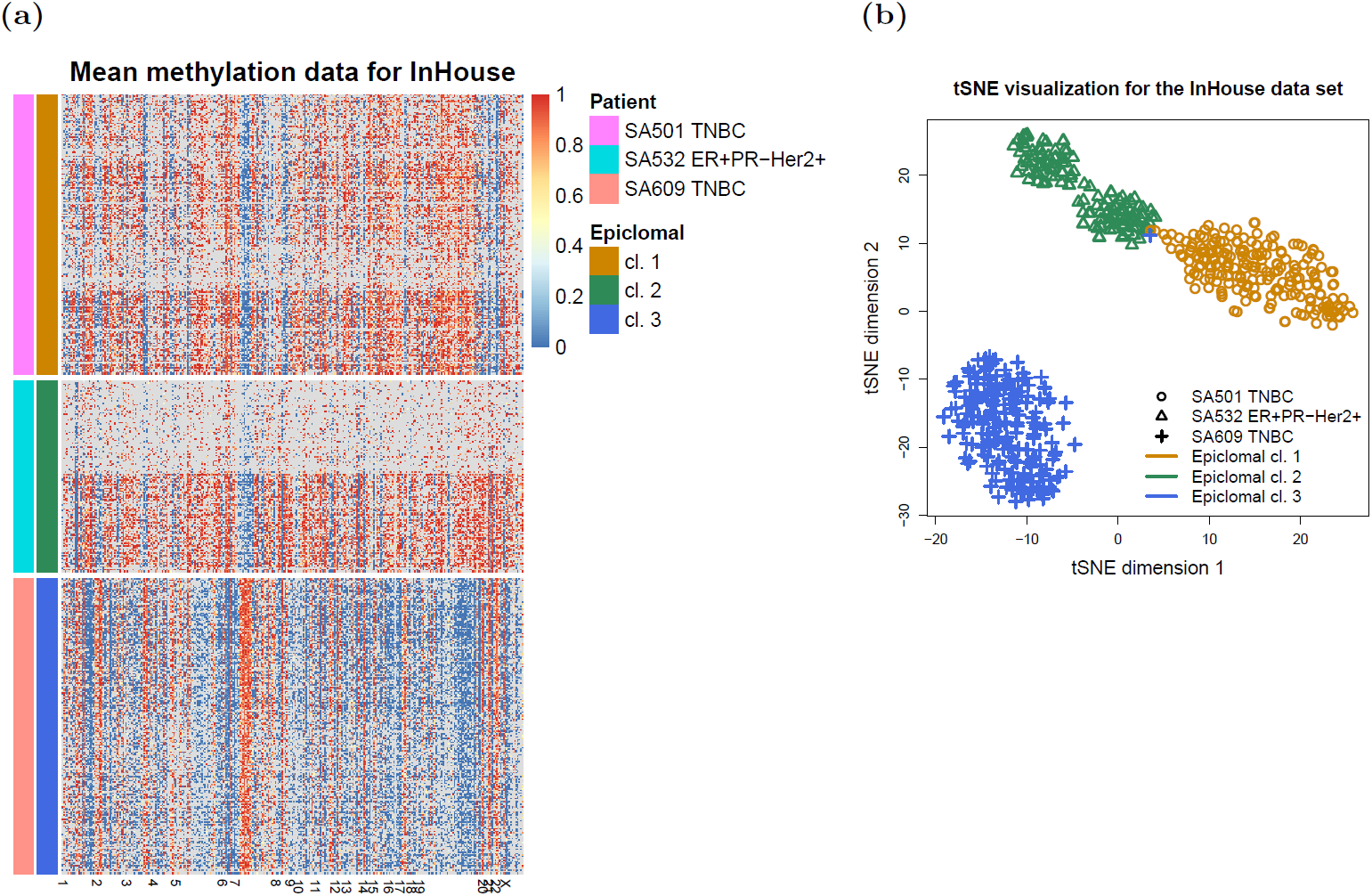
Visualization of the InHouse clusters. (a) EpiclomalRegion clustering, data filtered to include the most variable CGIs and obtain ≈ 15 000 loci (327 CGIs, cell average missing proportion 0.82, 558 cells). EpiclomalRegion obtained 3 clusters, V-measure = 1. Rows are cells and columns are CGIs. (b) tSNE dimensionality reduction and color-coding of the Epiclomal clusters onto the tSNE 2-dimensional space.

We next focused our analysis on one of the three patient-derived xenografts (PDX) above, which was previously characterized with whole-genome sequencing (WGS) [23] and single-cell WGS [4] (patient SA501 in Supplementary Table 1). Breast cancers often exhibit whole chromosome gains and losses (in addition to sub-chromosomal aneuploidy), especially of the X chromosome, which provides a strong methylation signal. As previously described, this PDX undergoes copy number clonal dynamics between passages, with clones losing one copy of X eventually dominating the populations of later passages. Patient tumour cells at diagnosis were mouse xenografted and serially transplanted over generations. Then, sc-WGBS data from passages 2, 7 and 10 were generated using the PBAL protocol [7]. After filtering out cells that did not pass quality control upon alignment (see Methods), we obtained a final sc-WGBS dataset of 244 single cells over 3 passages. We considered as initial regions the set of differentially methylated CGIs found when comparing bulk BS-seq data from passages 1 and 10 (see Methods). We then applied non-negative matrix factorization (NMF - [15, 17, 30]) to the region mean methylation data of all 244 cells as a feature selection strategy obtaining a final input set of 94 regions (see Supplementary Table 2 for their coordinates). Over all 94 regions (Supplementary Figures 14a and 14b) chromosome X contained the most differentially methylated regions of any single chromosome (29 out of 94; Supplementary Figure 13).

Using these 94 regions, EpiclomalRegion clustered the cells (Figure 7a) into four epiclones: two primarily containing passage 2 cells, and two containing a mix of passage 7 and 10 cells (EpiclomalBasic produced the same results). The distribution of posterior cluster assignment probabilities (*p*) indicates most cells were classified with *p*>0.9, with the exception of two cells that were assigned to Cluster 3 with probabilities of 0.73 and 0.69.

**Fig 7.**
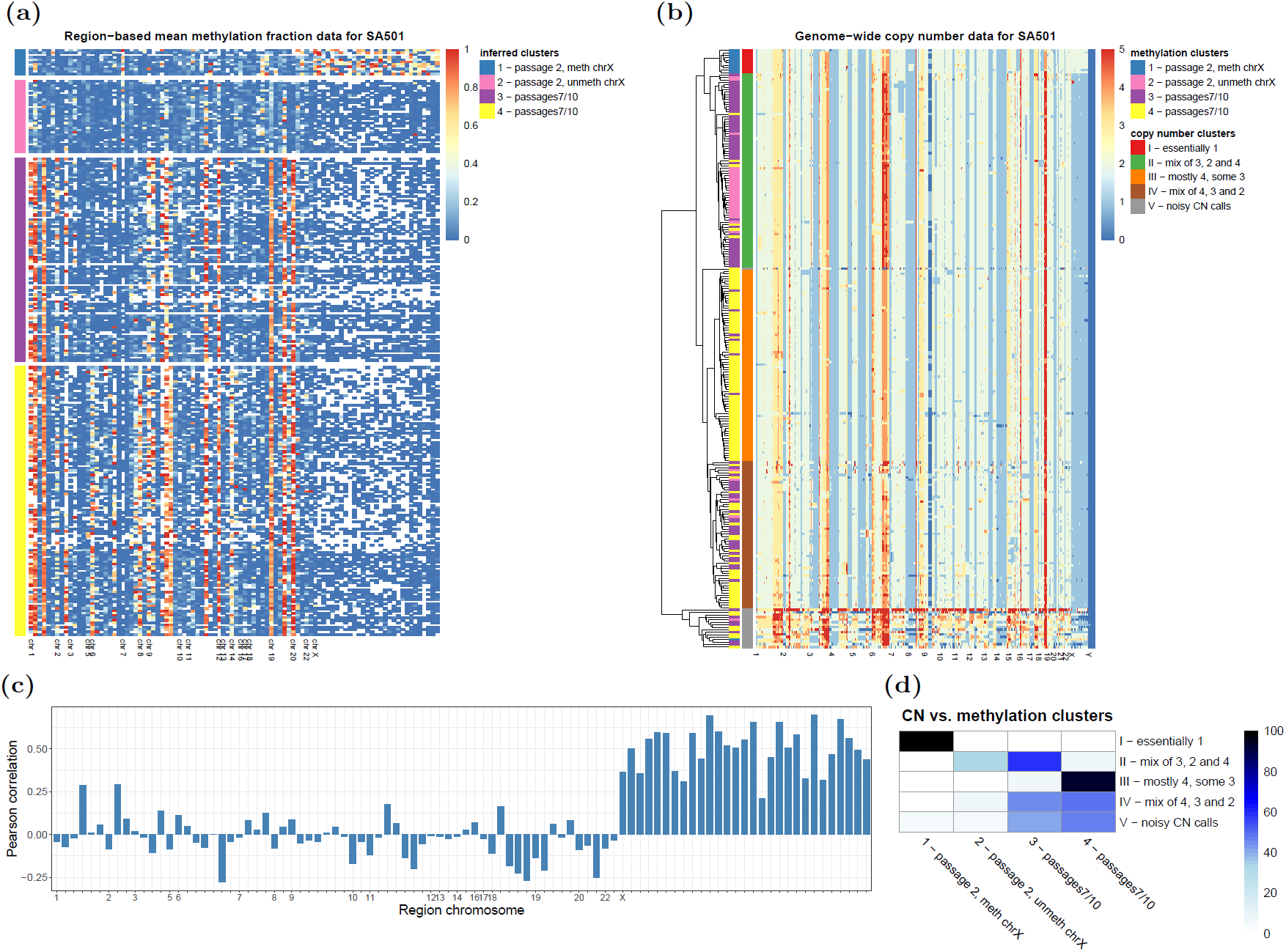
Results for patient SA501. (a) Mean methylation level for each of the 94 NMF-selected regions (CGIs) for patient SA501 across all cells ordered according to the four methylation clusters found using EpiclomalRegion. Rows are cells and columns are CGIs. (b) Inferred genome-wide copy numbers for the same cells as in (a) clustered using a ward.D2 hierarchical clustering method and Euclidean copy number distances. Note that copy number 5 actually means 5 or more copies. To call copy number changes, we used the methylation sc-WGBS data. Only one epiclone and one copy number clone match, the remaining clones transcend each other. (c) Pearson correlation between mean methylation data and copy number data in each of the 94 regions. There is correlation in chromosome X, but not in the autosomal chromosomes. (d) Heatmap showing the percentage of cells in the copy number clusters (rows) that are in the methylation clusters (columns); rows sum up to 100.

Inspection shows that Cluster 1 contains 10/40 passage 2 cells (and 1 passage 7 cell) with unmethylated features, except for chromosome X regions, which are mainly methylated. Cluster 2 contains the remaining 30/40 passage 2 cells (and one passage 10 cell), but with unmethylated chromosome X regions, consistent with X inactivation partitioning of the X chromosomes. At later passages, several autosomal regions became methylated (see, for example, chromosomes 1, 9, 12, 19 and 20, Supplementary Table 3). In addition, we identified three main regions that are methylated only in some of the later passage cells (see also Supplementary Table 3), resulting in two different epiclones each containing a mix of passage 7 and 10 cells (cluster 3 containing 53/98 passage 7 cells and 34/106 passage 10 cells; and cluster 4 containing 44/98 passage 7 cells and 71/106 passage 10 cells).

The observations above suggest that some chromosomal regions, such as X, may show strong copy number influence on CpG states, whereas others may differ in CpG state, but unrelated to copy number state in aneuploid genomes. Therefore we next investigated possible correlations between methylation and copy number alterations derived from the same sc-WGBS data (see Methods). A systematic comparison shows that indeed the average methylation levels and copy number states across cells for each of the 94 regions (Figure 7b) are only highly correlated (Pearson correlation >0.5; Figure 7c) for the X chromosome. This implies that epiclones may transcend copy number defined clones.

Indeed, when we compared the four epiclones with the four sc-PBAL copy number (CN) clones we noticed that they can match or transcend each other as follows (Figure 7b and d): epiclone 1 with methylated regions in the X chromosome matches exactly CN clone I having two copies of the X chromosome, which shows a strong relationship between the presence of the second copy of the X chromosome and the methylation pattern. However, 26/31 passage-2 cells with all 94 regions unmethylated from epiclone 2 are found in CN clone II, which also contains 47/87 cells from epiclone 3 and 6/115 cells from epiclone 4, even though these have several regions that are methylated. Finally, epiclone 3 transcends CN clones II (47/87 cells from epiclone 3 are in CN clone II) and IV (27/87 cells from epiclone 3 are in CN clone IV); and epiclone 4 transcends CN clones III (72/115 cells from epiclone 4 are in clone III) and IV (29/115 cells from epiclone 4 are in CN clone IV). Taken together, these data show for the first time with single cell methylation analysis that epigenetically defined clones may present a different lineage to that of copy number defined clonal architecture, opening up this form of analysis for cancer genomes.

## Discussion

Single cell CpG genome analysis is currently held back by a dearth of principled methods for handling the features of single cell methylation data. To this end, we have developed Epiclomal, a probabilistic CpG-based clustering method for clustering sparse sc-WGBS data and elucidation of epigenetic diversity on different types of cell populations. Our method has produced overall better results than non-probabilistic based methods when tested on synthetic data from seven extensive simulation scenarios (Figures 3 and 4) and four comprehensive real data sets (Figure 5). Epiclomal can impute the missing CpG methylation values more correctly than a naive imputation for the same clustering result (see panel h of Supplementary Figures 2-8). Importantly, Epiclomal is robust and reliable when the amount of data missing is large and/or varies across cells, and can find the true clusters and epiclone prevalence when the signal is subtle, both limiting features of current sc-WGBS data.

It is well understood that 5mC distribution in the genome is regionally clustered and this has implications for computational methods. EpiclomalRegion considers CpG-based methylation dependencies in functional regions and models errors while simultaneously assigning cells to clusters and imputing the missing data. It can also use bulk DNA methylation data to improve the epiclone prevalences, an important measure particularly to the study of cancer tumour composition. Epiclomal works at the CpG level and thus considers the contribution of every sequenced CpG site in the selected regions, without the loss of information by region averaging. Epiclomal does not only run an uninformed clustering method, but uses the clustering results of four other methods (with more easily added) and a robust model selection strategy to return the best prediction.

Epiclomal is part of an extensive statistical and computational framework that provides interpretable results and five different performance measures. It includes a pre-processing step where specific regions can be selected in order to increase signal and eliminate noise in the input data. Our framework also allows for novel components to be easily included in the computational pipeline. The non-probabilistic methods obtained better results, for example, on the smaller filtered Smallwood2014 input data sets than on the large input, supporting the notion that filtering out the most invariant regions may improve signal for clustering. In addition, our synthetic experiments as well as the SA501 intrapatient analysis on a well-designed set of differentially methylated regions showed that pre-processing the initial whole-genome data set in a way that keeps the clone differences and eliminates noise is likely to produce better results overall. Our selection strategy has the limitation of possibly removing regions that vary only in a small percentage of cells, which may result in clusters being condensed together. Future work includes a region selection strategy that can increase the signal-to-noise ratio. One approach, for example, would be to consider the variation across sites within regions so that regions with same variation pattern across cells could be represented only once in the model via appropriate weights in the data log-likelihood.

Although epigenomic states are of importance in cancer biology, to date very few single-cell whole genome bisulfite datasets have been generated on aneuploid cancer genomes. Here we used Epiclomal with a large (598 genomes) new dataset of sc-WGBS generated by the recently published PBAL method, to demonstrate how epiclones and copy number determined clones differ. Epiclomal was able to identify known and novel CpG methylation substructure that could not be identified by non-probabilistic distance based methods, due to the missing data inherent in sc-WGBS. Specifically, the separation between the two passage 7/10 subclusters, was not found by any of the non-probabilistic methods we considered, nor when a larger set of regions was used. This demonstrates that sophisticated modeling of the data missingness and appropriate region selection are necessary in order to clearly separate biological signal when this signal is not so abundant.

The ability to identify CpG defined sub-clones, epiclones, allowed us for the first time to compare a copy number determined lineage with an epigenetically defined lineage. It is expected that for certain regions of the genome, for example where allelic hemi-methylation occurs, changes in chromosomal copy number would strongly pattern 5mC CpG status. Indeed, we were able to observe this with subclones of a breast cancer PDX (SA501) where biallelic X chromosome clones present in early passages contain epiclones with and without CpG methylation, whereas for autosomes the copy number relationship is much weaker. In contrast, we observe that clones defined by autosomal copy number aberrations can exhibit quite distinct epiclone structure, leading to the notion that in some cases epiclone defined lineage will transcend that of copy number defined lineage. This has important implications for the study of cancer evolution and clonal states, as a failure to include epigenetic states will under-represent the cellular population structures of interest. Further work is required to define the scope and nature of epiclone versus copy number clone defined cellular lineages in cancer.

## Methods

### Pre-processing of the real data

We pre-process real data sets using the first part of our proposed framework. For each data set, we started considering all regions of corresponding type presented in the fifth column of Table 2. Then, after eliminating the empty regions across all cells, we also removed regions with an average missing proportion across all cells greater than or equal to 95%. Next, we kept the most variable regions (as measured by IQR of mean methylation levels) that would produce three filtered inputs with 10 000, 15 000 and 20 000 loci, respectively.

### In-house sc-WGBS data generation

#### Biospecimen collection and ethical approval

Tumour fragments from women diagnosed with breast lump undergoing surgery or diagnostic core biopsy were collected with informed consent, according to procedures approved by the Ethics Committees at the University of British Columbia. Patients in British Columbia were recruited and samples collected under tumor tissue repository (TTR-H06-00289) protocol that falls under UBC BCCA Research Ethics Board.

#### Tissue processing

The tumor materials were processed as mentioned in [23]. Briefly, the tumor fragments were minced finely with scalpels then mechanically disaggregated for one minute using a Stomacher 80 Biomaster (Seward Limited, Worthing, UK) in 1-2 mL cold DMEM-F12 medium. Aliquots from the resulting suspension of cells and clumps were used for xenotransplants.

#### Xenografting

Xenograft samples were transplanted and passaged as described in [23]. Female immune compromised, NOD/SCID interleukin-2 receptor gamma null (NSG) and NOD Rag-1 null interleukin-2 receptor gamma null (NRG) mice were bred and housed at the Animal Resource Centre (ARC) at the British Columbia Cancer Research Centre (BCCRC) supervised by Aparicio lab. Surgery was carried out on mice between the ages of 8-12 weeks. The animal care committee and animal welfare and ethical review committee, the University of British Columbia (UBC), approved all experimental procedures. For subcutaneous transplants, mechanically disaggregated cells and clumps of cells were resuspended in 100-200*µ*l of a 1:1 v/v mixture of cold DMEM/F12: Matrigel (BD Biosciences, San Jose, CA, USA). 8-12 weeks old mice were anesthetised with isoflurane, then the cell/clumps suspension was injected under the skin on the flank using a 1ml syringe and 21gauge needle.

#### Histopathological review

On histopathological review, two out of three, i.e. SA501 and SA609, patient derived xenografts (PDX) used in this study are triple negative breast cancers (TNBC). On immunohistochemistry, they were found to be receptor negative breast cancer subtype. SA532 is ER+PR-HER2+ xenografts. A pathologist reviewed the slides.

#### Cell preparation and dispensing

Xenograft tissues were dissociated to cells as described in [4] before dispensing single cells into the wells of 384 well plates using a contactless pieoelectric dispenser (sciFLEXArrayer S3, Scienion) with real-time cell detection in the glass capillary nozzle (CellenOne).

#### sc-WGBS experimental protocol

The Post-Bisulfite Adapter Ligation (PBAL) protocol described in [7] was used to obtain our in-house sc-WGBS data.

#### Data alignment and methylation calls

One lane of paired end sequencing was used to create each single cell library. Trim Galore (v0.4.1) and Cutadapt(v1.10) was used for quality and adapter trimming. Libraries were aligned to GRCh37-lite reference using Novoalign (v3.02.10) in bisulphite mode, and converted to BAM format and sorted using Sambamba (v0.6.0). Bam files were annotated for duplicates using Picard Tools’ MarkDuplicates.Jar (v1.92). Novomethyl (v1.10) was used in conjunction with in house scripts (samtools v1.6 and bedtools v2.25.0) to determine methylation of each CpG as described in Section “NovoMethyl - Analysing Methylation Status” Section of the Novoalign documentation (http://www.novocraft.com/documentation/novoalign-2/novoalign-user-guide/bisulphite-treated-reads/novomethyl-analyzing-methylation-status/).

#### Quality control

Using an in-house script, libraries were filtered according to a delta CT and 100K read count threshold to account for suitable library depth. Libraries over the expected number of CNV, were filtered out to control for chromothripsis and shattered cells.

#### Copy number calling

Copy number changes for SA501 were called using the same sc-WGBS DNA methylation data (copy number calling from the DLP protocol [4] largely match the sc-WGBS copy number calling for passage 2, results to appear). Control Free-c (v7.0) was used to copy number variant call on processed BAMs. The following settings were used: ploidy : 2, window and telocentromeric : 500000, sex : XY, minExpectGC : 0.39, and maxExpectedGC: 0.51.

### In-house bulk whole genome bisulfite sequencing (SA501 passages 1 and 10)

#### Whole genome bisulfite library construction for Illumina sequencing

To track the efficiency of bisulfite conversion, 10 ng lambda DNA (Promega) was spiked into 1 *µ*g genomic DNA quantified using Qubit fluorometry and arrayed in a 96-well microtitre plate. DNA was sheared to a target size of 300 bp using Covaris sonication and the fragments end repaired using DNA ligase and dNTPs at 30°C for 30 min. Repaired DNA was purified using a 2:1 AMPure XP beads to sample ratio and eluted in 40 *µ*L elution buffer in preparation for A-tailing; the addition of adenosine to the 3’ end of DNA fragments using Klenow fragment and dATP incubated at 37°C for 30 min.

Following reaction clean-up with magnetic beads, cytosine methylated paired-end adapters (5’-A^*m*^CA^*m*^CT^*m*^CTTT^*m*^C^*m*^C^*m*^CTA^*m*^CA^*m*^CGA^*m*^CG^*m*^CT^*m*^CTT^*m*^C^*m*^CGAT^*m*^CT-3’ and 3’-GAG^*m*^C^*m*^CGTAAGGA^*m*^CGA^*m*^CTTGG^*m*^CGAGAAGG^*m*^CTAG-5’) were ligated to the DNA at 30°C, 20 min and adapter flanked DNA fragments bead purified. Bisulfite conversion of the methylated adapter-ligated DNA fragments was achieved using the EZ Methylation-Gold kit (Zymo Research) following the manufacturer’s protocol. Seven cycles of PCR using HiFi polymerase (Kapa Biosystems) was used to enrich the bisulfite converted DNA and introduce fault tolerant hexamer barcode sequences. Post-PCR purification and size-selection of bisulfite converted DNA was performed using 1:1 AMPure XP beads. To determine final library concentrations, fragment sizes were assessed using a high sensitivity DNA assay (Agilent) and DNA quantified by Qubit fluorometry. Where necessary, libraries were diluted in elution buffer supplemented with 0.1% Tween-20 to achieve a concentration of 8nM for Illumina HiSeq2500 flowcell cluster generation.

#### Data alignment and methylation calls

FASTQ files were trimmed with TrimGalore (0.4.1) and then input into Bismark (0.14.4) aligning with bowtie2 (2.2.6). With the output BAM, we use samtools (1.3) to sort by name, fix mates, sort by position, remove duplicates, then finally sort by name once again and filter out reads with a mapping quality of 10 or less. We then ran the resulting BAM file through the bismark methylation extractor script that accompanies Bismark, to call methylation sites. All tools were run on all default settings, with changes made only to increase run speed.

#### Differentially methylated CpG Islands

Differentially methylated CpG islands between bulk samples from tumour xenograft passages 1 and 10 were obtained via Fisher’s exact test considering all CpG islands with coverage greater or equal than five reads. The Benjamini-Hochberg procedure was used to correct for multiple testing.

## Data Availability

Our raw InHouse and SA501 DNA methylation data has been uploaded to the European Genome-Phenome Archive (https://www.ebi.ac.uk/ega/home), accession number EGAS00001003504 and will be released upon acceptance for publication.

## Code Availability

Our computational code is available online at https://github.com/shahcompbio/Epiclomal.

## Author’s contributions

C.P.E.d.S. and M.A. developed and implemented the proposed methodology, performed all the analyses, and wrote the manuscript. T.M., F.K., J.B., E.M., J.B. and B.W. prepared the in-house single cells and bulk samples. Q.C. and M.W. generated the PBAL libraries. M.M. supervised library generation. R.M. supervised library sequencing. E.S., T.H. and D.L. processed the sequencing data and performed QC analyses. S.A. contributed to the manuscript text. M.H. and S.A. contributed ideas during the method development and analysis of the results. S.A. and S.P.S conceived and oversaw the project.

## Acknowledgements

We acknowledge generous funding support provided by the BC Cancer Foundation. SA is supported by grants from CIHR, Terry Fox Research Institute, Canadian Cancer Society Research Institute and the Breast Cancer Research Foundation. SPS receives operating funds from Terry Fox Research Institute (grant 1082) and the Canadian Cancer Society (grant 705636).

## References

1. Smith ZD, Meissner A. DNA methylation: roles in mammalian development. Nature Reviews Genetics. 2013;14(3):204.

2. Feng S, Jacobsen SE, Reik W. Epigenetic reprogramming in plant and animal development. Science. 2010;330(6004):622–627.

3. Navin N, Kendall J, Troge J, Andrews P, Rodgers L, McIndoo J, et al. Tumour evolution inferred by single-cell sequencing. Nature. 2011;472(7341):90.

4. Zahn H, Steif A, Laks E, Eirew P, VanInsberghe M, Shah SP, et al. Scalable whole-genome single-cell library preparation without preamplification. Nature methods. 2017;14(2):167.

5. Shapiro E, Biezuner T, Linnarsson S. Single-cell sequencing-based technologies will revolutionize whole-organism science. Nature Reviews Genetics. 2013;14(9):618.

6. Gawad C, Koh W, Quake SR. Single-cell genome sequencing: current state of the science. Nature Reviews Genetics. 2016;17(3):175.

7. Hui T, Cao Q, Wegrzyn-Woltosz J, O’Neill K, Hammond CA, Knapp DJHF, et al. High-Resolution Single-Cell DNA Methylation Measurements Reveal Epigenetically Distinct Hematopoietic Stem Cell Subpopulations. Stem Cell Reports. 2018;doi: https://doi.org/10.1016/j.stemcr.2018.07.003.

8. Clark SJ, Lee HJ, Smallwood SA, Kelsey G, Reik W. Single-cell epigenomics: powerful new methods for understanding gene regulation and cell identity. Genome biology. 2016;17(1):72.

9. Smallwood SA, Lee HJ, Angermueller C, Krueger F, Saadeh H, Peat J, et al. Single-cell genome-wide bisulfite sequencing for assessing epigenetic heterogeneity. Nature methods. 2014;11(8):817–820. February 21, 2020 12/23

10. Angermueller C, Clark SJ, Lee HJ, Macaulay IC, Teng MJ, Hu TX, et al. Parallel single-cell sequencing links transcriptional and epigenetic heterogeneity. Nature methods. 2016;13(3):229.

11. Farlik M, Halbritter F, Müller F, Choudry FA, Ebert P, Klughammer J, et al. DNA methylation dynamics of human hematopoietic stem cell differentiation. Cell stem cell. 2016;19(6):808–822.

12. Hou Y, Guo H, Cao C, Li X, Hu B, Zhu P, et al. Single-cell triple omics sequencing reveals genetic, epigenetic, and transcriptomic heterogeneity in hepatocellular carcinomas. Cell research. 2016;26(3):304.

13. Gravina S, Dong X, Yu B, Vijg J. Single-cell genome-wide bisulfite sequencing uncovers extensive heterogeneity in the mouse liver methylome. Genome biology. 2016;17(1):150.

14. Luo C, Keown CL, Kurihara L, Zhou J, He Y, Li J, et al. Single-cell methylomes identify neuronal subtypes and regulatory elements in mammalian cortex. Science. 2017;357(6351):600–604.

15. Mulqueen RM, Pokholok D, Norberg SJ, Torkenczy KA, Fields AJ, Sun D, et al. Highly scalable generation of DNA methylation profiles in single cells. Nature Biotechnology. 2018;.

16. Zeisel A, Muñoz-Manchado AB, Codeluppi S, Lönnerberg P, La Manno G, Juréus A, et al. Cell types in the mouse cortex and hippocampus revealed by single-cell RNA-seq. Science. 2015;347(6226):1138–1142.

17. Lee DD, Seung HS. Learning the parts of objects by non-negative matrix factorization. Nature. 1999;401(6755):788.

18. van der Maaten L, Hinton G. Visualizing Data using t-SNE. Journal of Machine Learning Research. 2008;9:2579–2605.

19. Ester M, Kriegel HP, Sander J, Xu X. A Density-Based Algorithm for Discovering Clusters in Large Spatial Databases with Noise. In: Proc. of 2nd International Conference on Knowledge Discovery and; 1996. p. 226–231.

20. Kapourani CA, Sanguinetti G. Melissa: Bayesian clustering and imputation of single cell methylomes. 2018;doi: http://dx.doi.org/10.1101/312025.

21. Angermueller C, Lee HJ, Reik W, Stegle O. DeepCpG: accurate prediction of single-cell DNA methylation states using deep learning. Genome biology. 2017;18(1):67.

22. DeRose YS, Wang G, Lin YC, Bernard PS, Buys SS, Ebbert MT, et al. Tumor grafts derived from women with breast cancer authentically reflect tumor pathology, growth, metastasis and disease outcomes. Nature medicine. 2011;17(11):1514.

23. Eirew P, Steif A, Khattra J, Ha G, Yap D, Farahani H, et al. Dynamics of genomic clones in breast cancer patient xenografts at single cell resolution. Nature. 2015;518(7539):422.

24. Spiegelhalter DJ, Best NG, Carlin BP, Van Der Linde A. Bayesian measures of model complexity and fit. Journal of the Royal Statistical Society: Series B (Statistical Methodology). 2002;64(4):583–639.

25. Nowell PC. The clonal evolution of tumor cell populations. Science. 1976;194(4260):23–28.

26. Ding J, Shah S, Condon A. densityCut: an efficient and versatile topological approach for automatic clustering of biological data. Bioinformatics. 2016;32(17):2567–2576.

27. Caliński T, Harabasz J. A dendrite method for cluster analysis. Communications in Statistics-theory and Methods. 1974;3(1):1–27.

28. Rosenberg A, Hirschberg J. V-measure: A conditional entropy-based external cluster evaluation measure. In: Proceedings of the 2007 joint conference on empirical methods in natural language processing and computational natural language learning (EMNLP-CoNLL); 2007.

29. Aryee M, Liu W, Engelmann J, Nuhn P, Gurel M, Haffner M, et al. DNA methylation alterations exhibit intraindividual stability and interindividual heterogeneity in prostate cancer metastases. Science Translational Medicine. 2013;5(169). doi:10.1126/scitranslmed.3005211.

30. Kim H, Park H. Sparse non-negative matrix factorizations via alternating non-negativity-constrained least squares for microarray data analysis. Bioinformatics. 2007;23(12):1495–1502.

