## Supplementary Material for "Epiclomal: probabilistic clustering of sparse single-cell DNA methylation data"

**5 Supplementary Tables** **15**

### 1 Non-probabilistic clustering methods

#### 1.1 EuclideanClust

EuclideanClust is a region-based method in which we first compute for each cell the mean methylation level of each region of interest. Because of the sparsity of the data, we cluster the cells taking as input data not the original matrix of mean methylation levels, but instead we apply complete-linkage hierarchical clustering on the symmetric matrix of Euclidean distances between every pair of cells with a dissimilarity matrix based also on Euclidean distances. EuclideanClust is similar to the approach used by Smallwood et al. [1] and Angermuller et al. [2], with the difference that the regions are defined differently in our case (functional genomic regions) versus Smallwood (sliding windows across the genome) and Angermuller et al. (gene bodies). We use the Calinski-Harabasz (CH) index [3] to automatically choose the number of clusters that best fit the data.

#### 1.2 DensityCut

As in EuclideanClust we first compute for each cell the mean methylation level of each region of interest. We then use principal component analysis as a dimensionality reduction technique considering a maximum of 20 first principal components, and apply DensityCut, a density based clustering algorithm proposed by [4], to the resulting principal component scores. This method is somewhat similar to the approach proposed by Mulqueen et al. [5], except that they used a different dimensionality reduction technique (NMF) and a different density-based clustering algorithm (DBSCAN).

#### 1.3 HammingClust

This method is a CpG-based method as we consider the data from all individual CpGs from all regions of interest to cluster the cells. Because of the sparsity of the data, similarly to EuclideanClust, clustering is done by first calculating Hamming distance based dissimilarities between each pair of cells and then applying Ward's linkage hierarchical clustering with Euclidean distances on the matrix of Hamming dissimilarities. HammingClust is essentially equivalent with the recent PDclust of Hui et al. [6]. As in EuclideanClust, the CH-index is used to select the optimal number of clusters.

#### 1.4 PearsonClust

PearsonClust is also a CpG-based approach similar to HammingClust, except that instead of Hamming and Euclidean distances it is based entirely on Pearson correlation, that is, we first compute the Pearson correlation between every pair of cells and then apply Ward's linkage hierarchical clustering with again a Pearson-based dissimilarity matrix on the initial correlation matrix. This method is equivalent to the approach used by Hou et al. [7]. The CH-index is also used for best clustering partition.

##### 1.5 Pre-imputation of missing values

Sometimes the input matrix to the non-probabilistic methods is too sparse and either the hierarchical clustering or the CH-index method for choosing the number of clusters will fail to produce results. For the synthetic data, we just report this as a failure in order to understand what characteristics of the input data set make this failure happen. However, for the real data sets, we try to run the hierarchical methods without pre-imputation, and, if they fail, we rerun them after pre-imputation, see the star-labelled runs in Figure 5 of the main manuscript.

The pre-imputation step is performed as follows. For the region-based methods, we impute a missing region value with the average across all cells for that region. For the CpG-based methods, since the CpG methylation value should be 0 or 1, we replace a missing CpG with the most common (median) CpG value observed across all cells for that CpG locus. If this median is 0.5, we use 0 instead.

#### 2 Proposed probabilistic method - Epiclomal

Our proposed methodology extends the approach of [8] to single-cell DNA methylation data. In what follows we describe our model and Variational Bayes (VB) inference technique for the case we call EpiclomalRegion, which is based on the assumption that the probability of a given locus being methylated depends on the genomic region that locus is located and that loci in the same genomic region share the same methylation probability. Our EpiclomalBasic approach is a special case of EpiclomalRegion obtained by assuming that all loci belong to one single region sharing the same probability of being methylated and, therefore, can be obtained by setting $R = 1$ in all calculations and steps below. See graphical models in Figure 1 of the main text.

##### 2.1 Model and inference

Let us consider a set of $R$ regions in the genome (e.g., CpG islands, gene bodies, etc). Let $X_{nrl}$ be the observed methylation status (or epigenotype) for cell $n$ at locus $l$ of region $r$, for $n = 1, \ldots, N$, $r = 1, \ldots, R$ and $l = 1, \ldots, L_r$. Our approach allows for the set of loci with observed data to vary across cells, but for simplicity we write our model and inference derivations assuming there are data for all loci in all cells, i.e., assuming complete data. Each $X_{nrl}$ takes value in $\mathcal{S} = \{\text{unmethylated, methylated}\}$ or simply $\mathcal{S} = \{0, 1\}$.

Let $\mathbf{X}_{nr} = (X_{nr1}, \ldots, X_{nrL_r})^T$ be the vector of observed data for region $r$ in cell $n$ and $\mathbf{X}_n = (\mathbf{X}_{nr}^T, \ldots, \mathbf{X}_{nR}^T)^T$ be the vector with all observed data for cell $n$. We assume that

- $\mathbf{X}_1, \ldots, \mathbf{X}_N$ are independent;
- $\mathbf{X}_{n1}, \ldots, \mathbf{X}_{nR}$ are independent for all $n$;

- Given a vector of true methylation states, $X_{nr1}, \ldots, X_{nrL_r}$ are independent with the distribution of $X_{nrl}$ depending on the true methylation state at locus $l$ of region $r$.

We assume that there are $K << N$ vectors of true hidden methylation states shared across the cells. Let $Z_n$ taking values in $\{1, \ldots, K\}$ be the hidden variable indicating the true cluster (epiclonal) population of cell $n$. We consider $Z_1, \ldots, Z_N$ independent with $P(Z_n = k) = \pi_k$ such that $\sum_{k=1}^{K} \pi_k = 1$. If $Z_n = k$ then the distribution of $\mathbf{X}_n$ depends on the $k$-th vector of true hidden epigenotypes $\mathbf{G}_k = (\mathbf{G}_{k1}^T, \ldots, \mathbf{G}_{kR}^T)^T$, where $\mathbf{G}_{kr} = (G_{kr1}, \ldots, G_{krL_r})^T$. We consider that

- $\mathbf{G}_1, \ldots, \mathbf{G}_K$ are independent;
- $\mathbf{G}_{k1}, \ldots, \mathbf{G}_{kR}$ are independent for all $k$;
- $G_{kr1}, \ldots, G_{krL_r}$ are independent with $P(G_{krl} = s) = \mu_{krs}$ such that $\sum_{s \in \mathcal{S}} \mu_{krs} = 1$, that is, $G_{krl}$ follows a categorical distribution with parameter set $\boldsymbol{\mu}_{kr} = \{\mu_{krs} : s \in \mathcal{S}\}$.

Therefore, given the true cluster assignment and the corresponding true hidden methylation states, the observed data $\mathbf{X}_{nr}$ are independent with $X_{nrl}$ following a categorical distribution with parameters depending on the hidden true state at locus $l$ of region $r$ for cluster population $k$, that is,

$$P(X_{nrl} = t | Z_n = k, G_{krl} = s) = \epsilon_{st} \text{ with } \sum_{t \in \mathcal{S}} \epsilon_{st} = 1. \quad (1)$$

We can also interpret the probability in (1) as a misclassification error, which in this context is related to sequencing error.

Let $\Theta$ be the set containing all the model parameters, i.e., $\Theta = \{\boldsymbol{\mu}, \boldsymbol{\epsilon}, \boldsymbol{\pi}\}$, where

- $\boldsymbol{\mu} = (\boldsymbol{\mu}_1^T, \ldots, \boldsymbol{\mu}_K^T)^T$ with $\boldsymbol{\mu}_k = (\boldsymbol{\mu}_{k1}^T, \ldots, \boldsymbol{\mu}_{kR}^T)^T$ and $\boldsymbol{\mu}_{kr} = \{\mu_{krs} : s \in \mathcal{S}\}$;
- $\boldsymbol{\epsilon} = \{\boldsymbol{\epsilon}_s : s \in \mathcal{S}\}$ with $\boldsymbol{\epsilon}_s = \{\epsilon_{st} : t \in \mathcal{S}\}$ and
- $\boldsymbol{\pi} = (\pi_1, \ldots, \pi_K)^T$.

In order to infer $\Theta$ and the hidden states $\mathbf{Z} = (Z_1, \ldots, Z_n)^T$ and $\mathbf{G} = \{\mathbf{G}_1, \ldots, \mathbf{G}_K\}$ we take on a Bayesian approach and consider the Variational Bayes (VB) algorithm ([9] and [10]) to approximate the posterior distribution

$$q(\mathbf{Z}, \mathbf{G}, \Theta) \equiv P(\mathbf{Z}, \mathbf{G}, \Theta | \mathbf{X}). \quad (2)$$

We consider the following prior distributions for the parameters in $\Theta$.

- $p(\boldsymbol{\mu}) = \prod_{k=1}^{K} p(\boldsymbol{\mu}_k) = \prod_{k=1}^{K} \prod_{r=1}^{R} p(\boldsymbol{\mu}_{kr})$, where $\boldsymbol{\mu}_{kr} \sim \text{Dirichlet}(\boldsymbol{\beta}^0)$

- $p(\boldsymbol{\epsilon}) = \prod_{s\in\mathcal{S}} p(\boldsymbol{\epsilon}_s)$, where $\boldsymbol{\epsilon}_s \sim \text{Dirichlet}(\boldsymbol{\gamma}_s^0)$
- $\boldsymbol{\pi} \sim \text{Dirichlet}(\boldsymbol{\alpha}^0)$

In what follows we describe the main steps of the VB algorithm we used to infer $\mathbf{Z}, \mathbf{G}$ and $\Theta$.

*Step 1. Posterior factorization*

We assume the following factorization of the posterior distribution in (2):

$$\begin{aligned} q(\mathbf{Z}, \mathbf{G}, \Theta) &\equiv q(\mathbf{Z})q(\mathbf{G})q(\boldsymbol{\mu})q(\boldsymbol{\epsilon})q(\boldsymbol{\pi}) \\ &= \Big[\prod_{n=1}^{N} q(Z_n)\Big]\Big[\prod_{k=1}^{K}\prod_{r=1}^{R}\prod_{l=1}^{L_r} q(G_{krl})\Big]\Big[\prod_{s\in\mathcal{S}} q(\boldsymbol{\epsilon}_s)\Big]\Big[\prod_{k=1}^{K}\prod_{r=1}^{R} q(\boldsymbol{\mu}_{kr})\Big]q(\boldsymbol{\pi}). \end{aligned} \tag{3}$$

*Step 2. Joint distribution of observed data, hidden variables and parameters*

Considering the assumptions previously made for the observed data, hidden variables and model parameters, we can write the logarithm of the joint distribution of $\mathbf{X}$, $\mathbf{Z}$, $\mathbf{G}$ and the parameters in $\Theta$ as

$$\log P(\mathbf{X}, \mathbf{Z}, \mathbf{G}, \Theta) = \log P(\mathbf{X}|\mathbf{Z}, \mathbf{G}, \Theta) + \log P(\mathbf{G}|\Theta) + \log P(\mathbf{Z}|\Theta) + \log P(\Theta), \tag{4}$$

where

$$\log P(\mathbf{X}|\mathbf{Z}, \mathbf{G}, \Theta) = \sum_{n=1}^{N}\sum_{r=1}^{R}\sum_{l=1}^{L_r}\sum_{k=1}^{K}\sum_{s\in\mathcal{S}}\sum_{t\in\mathrm{S}} \mathrm{I}(Z_n = k)\mathrm{I}(X_{nrl} = t)\mathrm{I}(G_{krl} = s)\log \epsilon_{st}; \tag{5}$$

$$\log P(\mathbf{G}|\Theta) = \sum_{k=1}^{K}\sum_{r=1}^{R}\sum_{l=1}^{L_r}\sum_{s\in\mathcal{S}} \mathrm{I}(G_{krl} = s)\log \mu_{krs}; \tag{6}$$

$$\log P(\mathbf{Z}|\Theta) = \sum_{n=1}^{N}\sum_{k=1}^{K} \mathrm{I}(Z_n = k)\log \pi_k \text{ and} \tag{7}$$

$$\log P(\Theta) = \log P(\boldsymbol{\epsilon}) + \log P(\boldsymbol{\mu}) + \log P(\boldsymbol{\pi})$$

with

$$\log P(\boldsymbol{\epsilon}) = \sum_{s\in\mathcal{S}}\Big[\sum_{t\in\mathcal{S}}(\gamma_{st}^0 - 1)\log \epsilon_{st} - \log B(\boldsymbol{\gamma}_s^0)\Big]; \tag{8}$$

$$\log P(\boldsymbol{\mu}) = \sum_{k=1}^{K}\sum_{r=1}^{R}\Big[\sum_{s\in\mathcal{S}}(\beta_s^0 - 1)\log \mu_{krs} - \log B(\boldsymbol{\beta}^0)\Big] \text{ and} \tag{9}$$

$$\log P(\boldsymbol{\pi}) \;=\; \sum_{k=1}^{K}(\alpha_k^0 - 1)\log \pi_k - \log B(\boldsymbol{\alpha}^0). \tag{10}$$

The function $B$ in (8)-(10) is the multivariate Beta function, which can be expressed in terms of gamma functions. So, for example,

$$B(\boldsymbol{\alpha}^0) = \frac{\prod_{k=1}^{K}\Gamma(\alpha_k^0)}{\Gamma(\sum_{k=1}^{K}\alpha_k^0)}.$$

*Step 3. Approximation*

We now approximate each term in the factorization (3) by calculating the expectation of $\log P(\mathbf{X}, \mathbf{Z}, \mathbf{G}, \Theta)$ over the distribution of all terms except the one of interest. So, for example, we obtain the approximation $q^*(\boldsymbol{\pi})$ for $q(\boldsymbol{\pi})$ by calculating

$$\log q^*(\boldsymbol{\pi}) = \mathbb{E}_{\mathbf{Z},\mathbf{G},\boldsymbol{\epsilon},\boldsymbol{\mu}}\left(\log P(\mathbf{X}, \mathbf{Z}, \mathbf{G}, \Theta)\right) + C, \tag{11}$$

where the expectation is taken with respect to the posterior distributions of $\mathbf{Z}$, $\mathbf{G}$, $\boldsymbol{\epsilon}$ and $\boldsymbol{\mu}$. This form of approximation arises from finding the distribution that minimizes the Kullback-Leibler divergence to the exact posterior, which is equivalent to maximizing the evidence lower bound (ELBO) given by

$$\text{ELBO}(q) = \mathbb{E}\left[\log P(\mathbf{X}, \mathbf{Z}, \mathbf{G}, \Theta)\right] - \mathbb{E}\left[\log q(\mathbf{Z}, \mathbf{G}, \Theta)\right]. \tag{12}$$

See [9] and [10] for more details.

In what follows we show how we obtain $q^*$ for each of our quantities of interest.

- Approximating $q(\boldsymbol{\pi})$ by $q^*(\boldsymbol{\pi})$

  We find $q^*(\boldsymbol{\pi})$ that satisfies

$$\log q^*(\boldsymbol{\pi}) = \mathbb{E}_{\mathbf{Z},\mathbf{G},\boldsymbol{\epsilon},\boldsymbol{\mu}}\left(\log P(\mathbf{X}, \mathbf{Z}, \mathbf{G}, \Theta)\right) + C.$$

  As only the terms (7) and (10) in (4) depend on $\boldsymbol{\pi}$ we calculate:

$$\begin{aligned}
\log q^*(\boldsymbol{\pi}) \;&=\; \mathbb{E}_{\mathbf{Z},\mathbf{G},\boldsymbol{\epsilon},\boldsymbol{\mu}}\left(\log P(\mathbf{Z}|\Theta)\right) \;+\; \mathbb{E}_{\mathbf{Z},\mathbf{G},\boldsymbol{\epsilon},\boldsymbol{\mu}}\left(\log P(\boldsymbol{\pi})\right) \;+\; C^* \\
&=\; \sum_{n=1}^{N}\sum_{k=1}^{K}\mathbb{E}_{q^*(Z_n)}\left(\mathrm{I}(Z_n = k)\right)\log \pi_k + \log P(\boldsymbol{\pi}) + C^* \\
&=\; \sum_{k=1}^{K}\log \pi_k\Big[\sum_{n=1}^{N}\mathbb{E}_{q^*(Z_n)}\left(\mathrm{I}(Z_n = k)\right)\Big] + \sum_{k=1}^{K}\log \pi_k(\alpha_k^0 - 1) + C^{**} \\
&=\; \sum_{k=1}^{K}\log \pi_k\Big[\Big(\sum_{n=1}^{N}\mathbb{E}_{q^*(Z_n)}\left(\mathrm{I}(Z_n = k)\right) + \alpha_k^0\Big) - 1\Big] + C^{**}.
\end{aligned} \tag{13}$$

Therefore, $q^*(\boldsymbol{\pi})$ is a Dirichlet distribution with parameters $\alpha^* = (\alpha_1^*, \ldots, \alpha_K^*)^T$, where

$$\alpha_k^* = \alpha_k^0 + \sum_{n=1}^{N} \mathbb{E}_{q^*(Z_n)} \big(\mathrm{I}(Z_n = k)\big). \tag{14}$$

- Approximating $q(Z_n)$ by $q^*(Z_n)$

  To find $q^*(Z_n)$ we calculate

$$\log q^*(Z_n) = \mathbb{E}_{Z_{i:i\neq n}, \mathbf{G}, \boldsymbol{\epsilon}, \boldsymbol{\mu}} \big(\log P(\mathbf{X}, \mathbf{Z}, \mathbf{G}, \Theta)\big) + C. \tag{15}$$

As only (5) and (7) in (4) depend on $Z_n$, calculating the approximation in (15) is equivalent to calculating the following:

$$\log q^*(Z_n) = \mathbb{E}_{Z_{i:i\neq n}, \mathbf{G}, \boldsymbol{\epsilon}, \boldsymbol{\mu}} \big(\log P(\mathbf{X}|\mathbf{Z}, \mathbf{G}, \Theta)\big) + \mathbb{E}_{Z_{i:i\neq n}, \mathbf{G}, \boldsymbol{\epsilon}, \boldsymbol{\mu}} \big(\log P(\mathbf{Z}|\Theta).\big) \tag{16}$$

Note that we can split $\log P(\mathbf{X}|\mathbf{Z}, \mathbf{G}, \Theta)$ in (5) into two terms: one that depends on $Z_n$ and one that is constant on $Z_n$, that is,

$$\begin{aligned} \log P(\mathbf{X}|\mathbf{Z}, \mathbf{G}, \Theta) &= \sum_{r=1}^{R} \sum_{l=1}^{L_r} \sum_{k=1}^{K} \sum_{s\in\mathcal{S}} \sum_{t\in\mathrm{S}} \mathrm{I}(Z_n = k)\mathrm{I}(X_{nrl} = t)\mathrm{I}(G_{krl} = s) \log \epsilon_{st} \\ &+ \sum_{i\neq n} \sum_{r=1}^{R} \sum_{l=1}^{L_r} \sum_{k=1}^{K} \sum_{s\in\mathcal{S}} \sum_{t\in\mathrm{S}} \mathrm{I}(Z_i = k)\mathrm{I}(X_{irl} = t)\mathrm{I}(G_{krl} = s) \log \epsilon_{st}. \end{aligned} \tag{17}$$

Similarly, we can write

$$\begin{aligned} \log P(\mathbf{Z}|\Theta) &= \sum_{k=1}^{K} \mathrm{I}(Z_n = k) \log \pi_k \\ &+ \sum_{i\neq n} \sum_{k=1}^{K} \mathrm{I}(Z_i = k) \log \pi_k. \end{aligned} \tag{18}$$

Therefore, considering the second terms in (17) and (18) as constants and taking the expectation in (16), we obtain:

$$\begin{aligned} \log q^*(Z_n) &= \sum_{k=1}^{K} \mathrm{I}(Z_n = k) \Big\{ \mathbb{E}_{q^*(\boldsymbol{\pi})}(\log \pi_k) \\ &+ \sum_{r=1}^{R} \sum_{l=1}^{L_r} \sum_{s\in\mathcal{S}} \sum_{t\in\mathcal{S}} \mathbb{E}_{q^*(G_{krl})} \big(\mathrm{I}(G_{krl} = s)\big)\mathrm{I}(X_{nrl} = t)\, \mathbb{E}_{q^*(\boldsymbol{\epsilon}_s)}(\log \epsilon_{st}) \Big\} + C^{**}. \end{aligned}$$

So that $q^*(Z_n) \sim \text{Categorical}(\boldsymbol{\pi}_n^*)$ with parameters $\boldsymbol{\pi}_n^* = (\pi_{n1}^*, \ldots, \pi_{nK}^*)^T$ where $\sum_{k=1}^{K} \pi_{nk}^* = 1$ and each $\pi_{nk}^*$ is given by

$$\pi_{nk}^* = \frac{\exp\left\{ \mathbb{E}_{q^*(\boldsymbol{\pi})}(\log \pi_k) + \sum_{r=1}^{R} \sum_{l=1}^{L_r} \sum_{s \in \mathcal{S}} \sum_{t \in \mathcal{S}} \mathbb{E}_{q^*(G_{krl})} \big(\mathrm{I}(G_{krl} = s)\big) \mathrm{I}(X_{nrl} = t)\, \mathbb{E}_{q^*(\boldsymbol{\epsilon}_s)}(\log \epsilon_{st}) \right\}}{\sum_{j=1}^{K} \exp\left\{ \mathbb{E}_{q^*(\boldsymbol{\pi})}(\log \pi_j) + \sum_{r=1}^{R} \sum_{l=1}^{L_r} \sum_{s \in \mathcal{S}} \sum_{t \in \mathcal{S}} \mathbb{E}_{q^*(G_{jrl})} \big(\mathrm{I}(G_{jrl} = s)\big) \mathrm{I}(X_{nrl} = t)\, \mathbb{E}_{q^*(\boldsymbol{\epsilon}_s)}(\log \epsilon_{st}) \right\}}. \tag{19}$$

Using similar calculations as the ones above for $q^*(Z_n)$ and $q^*(\boldsymbol{\pi})$, we obtain the following posterior approximations for the remaining quantities of interest.

- $q^*(\boldsymbol{\mu}_{kr}) \sim \text{Dirichlet}(\boldsymbol{\beta}_{kr}^*)$ where $\boldsymbol{\beta}_{kr}^*$ is the vector containing $\beta_{krs}^*$ for every $s \in \mathcal{S}$ with

$$\beta_{krs}^* = \beta_s^0 + \sum_{l=1}^{L_r} \mathbb{E}_{q^*(G_{krl})} \big(\mathrm{I}(G_{krl} = s)\big) \;\text{ for all }\; s \in \mathcal{S}. \tag{20}$$

- $q^*(G_{krl}) \sim \text{Categorical}(\boldsymbol{\mu}_{krl}^*)$ where $\boldsymbol{\mu}_{krl}^* = \{\mu_{krls}^* : s \in \mathcal{S}\}$ with

$$\mu_{krls}^* = \frac{\exp\left\{ \sum_{n=1}^{N} \sum_{t \in \mathcal{S}} \mathbb{E}_{q^*(Z_n)} \big(\mathrm{I}(Z_n = k)\big) \mathrm{I}(X_{nrl} = t)\, \mathbb{E}_{q^*(\boldsymbol{\epsilon}_s)}(\log \epsilon_{st}) + \mathbb{E}_{q^*(\boldsymbol{\mu}_{kr})}(\log \mu_{krs}) \right\}}{\sum_{v \in \mathcal{S}} \exp\left\{ \sum_{n=1}^{N} \sum_{t \in \mathcal{S}} \mathbb{E}_{q^*(Z_n)} \big(\mathrm{I}(Z_n = k)\big) \mathrm{I}(X_{nrl} = t)\, \mathbb{E}_{q^*(\boldsymbol{\epsilon}_v)}(\log \epsilon_{vt}) + \mathbb{E}_{q^*(\boldsymbol{\mu}_{kr})}(\log \mu_{krs}) \right\}} \tag{21}$$

- $q^*(\boldsymbol{\epsilon}_s) \sim \text{Dirichlet}(\boldsymbol{\gamma}_s^*)$ where $\boldsymbol{\gamma}_s^* = \{\gamma_{st}^* : t \in \mathcal{S}\}$, where

$$\gamma_{st}^* = \gamma_{st}^0 + \sum_{n=1}^{N} \sum_{r=1}^{R} \sum_{l=1}^{L_r} \sum_{k=1}^{K} \mathbb{E}_{q^*(Z_n)} \big(\mathrm{I}(Z_n = k)\big)\, \mathbb{E}_{q^*(G_{krl})} \big(\mathrm{I}(G_{krl} = s)\big) \mathrm{I}(X_{nrl} = t) \tag{22}$$

*Step 4. Expectations and updates*

Let $\psi$ be the digamma function defined as

$$\psi(x) = \frac{d}{dx} \log \Gamma(x), \tag{23}$$

which can be easily calculated via numerical approximation. The values of the expectations in (19)-(22) taken with respect to the approximated distributions are given as follows.

$$\mathbb{E}_{q^*(Z_n)} \big(\mathrm{I}(Z_n = k)\big) = \pi_{nk}^*$$

$$\mathbb{E}_{q^*(G_{krl})}\big(\mathrm{I}(G_{krl}=s)\big) = \mu^*_{krls}$$
$$\mathbb{E}_{q^*(\boldsymbol{\epsilon}_s)} = \psi(\gamma^*_{st}) - \psi\Big(\sum_{t\in\mathcal{S}}\gamma^*_{st}\Big) \tag{24}$$

$$\mathbb{E}_{q^*(\boldsymbol{\mu}_{kr})}(\log\mu_{krs}) = \psi(\beta^*_{krs}) - \psi\Big(\sum_{s\in\mathcal{S}}\beta^*_{krs}\Big) \tag{25}$$

$$\mathbb{E}_{q^*(\boldsymbol{\pi})}(\log\pi_k) = \psi(\alpha^*_k) - \psi\Big(\sum_{k=1}^{K}\alpha^*_k\Big) \tag{26}$$

Using the results above regarding the expectations, we update the parameters of the approximated distributions iteratively as follows.

We initialize $\boldsymbol{\gamma}^*_s$, $\boldsymbol{\beta}^*_{kr}$ and $\boldsymbol{\pi}^*_n$ with some arbitrary values (e.g., $\boldsymbol{\gamma}^{*(0)}_s = \boldsymbol{\gamma}^0_s$ and $\boldsymbol{\beta}^{*(0)}_{kr} = \boldsymbol{\beta}^0$). At each iteration $c$ we compute:

1. $\boldsymbol{\alpha}^{*(c)}$ using $\boldsymbol{\pi}^{*(c-1)}_n$
2. $\boldsymbol{\mu}^{*(c)}_{krl}$ using $\boldsymbol{\gamma}^{*(c-1)}_s$, $\boldsymbol{\pi}^{*(c-1)}_n$ and $\boldsymbol{\beta}^{*(c-1)}_{kr}$
3. $\boldsymbol{\beta}^{*(c)}_{kr}$ using $\boldsymbol{\mu}^{*(c)}_{krl}$
4. $\boldsymbol{\gamma}^{*(c)}_s$ using $\boldsymbol{\mu}^{*(c)}_{krl}$ and $\boldsymbol{\pi}^{*(c-1)}_n$
5. $\boldsymbol{\pi}^{*(c)}_n$ using $\boldsymbol{\mu}^{*(c)}_{krl}$, $\boldsymbol{\gamma}^{*(c)}_s$ and $\boldsymbol{\alpha}^{*(c)}$

We then conduct many iterations of 1-5 until the convergence of the ELBO in (12).

#### 2.2 Initialization and choice of $K$

We run the Variational Bayes algorithm described in the previous section a maximum number times $T$ (we used $T = 1000$ for the real data sets and $T = 300$ for the synthetic data sets). We start from different initial $\boldsymbol{\pi}^*_n$ values for each cell $n$ (the other two values are initialized with the corresponding hyperparameter $\boldsymbol{\gamma}^{*(0)}_s = \boldsymbol{\gamma}^0_s$ and $\boldsymbol{\beta}^{*(0)}_{kr} = \boldsymbol{\beta}^0$). That is, each vector $\boldsymbol{\pi}^*_n$ of length $K$ will have $K-1$ values of 0 and one value of 1, corresponding to the initial cluster for that cell. Most initializations are uniformly random, but informative starting values often lead to better results. Therefore, for all analyses we use the following initialization strategy. First we run EuclideanClust and if the hierarchical clustering is successful we cut the hierarchical tree at $1, 2 \ldots K$ clusters, obtaining the first $K$ initial points. Then, we do the same for HammingClust and PearsonClust, obtaining $2 \times K$ more initial points. Finally, we add the prediction made by DensityCut. Note that initializations from more additional clustering methods can be easily added to our framework.

In our analyses we use $K = 10$ for all synthetic and real data sets. Therefore, a maximum of $I = 31$ initializations are from the non-probabilistic methods. The remaining $T - I$ VB runs are initialized randomly, with each initial number of clusters being a number chosen uniformly at random between 1 and $K$. For each run, the VB algorithm returns a number of recommended clusters $c \leq K$ and the corresponding cell-to-cluster assignments. With this strategy we obtain a more uniform number of clusters across all runs than if we use the same $K$ for each run. Therefore, our strategy is similar to a BIC or AIC selection criterion in which we would perform a roughly equal number of runs for each possible number of recommended clusters.

After obtaining the $T$ runs (this was done in parallel on a computing cluster), we have for each run the number of recommended clusters $c \leq K$ and the computed DIC score that takes into account the likelihood of the model as well as the model complexity [11]. Then, for each $c$, we compute the minimum DIC obtained for all runs that recommended $c$ clusters, and we plot the DIC curve, such as the one in Supplementary Figure 1.

Now, having a DIC curve, we find the elbow point as follows. We draw a line from the first to the last point of the curve and then find the DIC point that is the farthest away from that line. Sometimes, the DIC curve is not a smooth decreasing function, but instead it can increase and decrease. Therefore, we have two versions of the DIC-elbow selection strategy:

1. A course-grained DIC-elbow selection strategy in which we consider only the part of the curve with DIC values decreasing by at least a small percentage threshold (for example 0.2%) - the green line in Supplementary Figure 1. We then find the elbow for this part of curve, which corresponds to the best choice of number of clusters - the red line in Supplementary Figure 1. This strategy is best used in cases when we are not looking for very minor clusters, but mostly for major clusters.

2. A fine-grained DIC-elbow selection strategy in which there is no threshold for the green line, in other words the green line is at the far right of the plot in Supplementary Figure 1. This strategy is best used when we are looking for minor clusters.

The DIC-elbow selection strategy can be seen as an automatic way to select the best run. However, visual inspection of the DIC-elbow can sometimes help choose the best thresholds.

##### 2.3 EpiclomalBulk

Often, bulk CpG-level methylation data are produced, that is, a vector of natural numbers, representing the number of methylated cytosines for each CpG, from 0 to the read depth $D$ (e.g., $D = 60$). For instance, a number of 0 means that we expect no cell to be methylated (all are unmethylated) at that CpG site. A number of 60 means that we expect all the cells to be methylated, and a number of

30 means that roughly half of the cells are methylated and half are unmethylated. Therefore, given the cell-to-cluster assignments and the corresponding imputed methylation values, we can compute a score that tells us how well the given imputed values match the bulk data (for each CpG site, we just have to count the number of cells that are methylated and then divide by the number of cells and multiply by $D$).

Having this bulk-based score function, we designed a stochastic local search algorithm that starts from a given configuration (this is EpiclomalRegion's best result), keeps the number of clusters fixed, and randomly reassigns "uncertain cells" to one of their "candidate clusters". The "uncertain cells" and the "candidate clusters" are obtained as described in Section 2.4. Only the CpGs in the regions that make the clusters different are considered. If the new score is better than before, we always keep it, if it is not, we only keep it 20% of the times to help the algorithm escape local minima. We repeat this strategy for 10 iterations and return the new cell-to-cluster assignments and imputed methylation states that gives the best score.

At the time of this publication the EpiclomalBulk feature has been implemented and tested on synthetic data with up to three correct number of clusters.

#### 2.4 Adjustment of imputed CpG states and naive imputation

In the cases when there are no observed values at a given CpG position for all cells in an estimated cluster, our probablistic model fills this value with the value that is the most common in the region of that CpG in that cluster. However, if this region is not a differentially methylated region, it may be better to just fill in with the CpG values from other clusters. Incorporating this mechanism in the Epiclomal model would require having another node influencing node $G$ that depends on each CpG position, along with a trade-off mechanism that considers both the influence of other CpGs in the region for the cluster, and the influence of the observed values for the same CpG in other clusters. We reasoned that this would be an unnecessarily complicated and expensive addition to our model. Therefore, we added a post-processing "imputation adjustment" step in which we go through every region, and, if the prediction implies that this region is differentially methylated, we do not change anything; otherwise, for all the imputed CpGs that were completely missing in the cluster, we use the median value of all cells or 0 (unmethylated) if this median is 0.5 or undefined. We call a region differentially methylated if there is a pair of clusters for which the $G$ vectors for the region have Pearson correlation coefficient less than 0.5. Panels h of Supplementart Figures 2-8 use this adjustment strategy for the EpiclomalRegion-adjusted case.

In order to assess Epiclomal's imputation of the missing CpG states, we also calculate a naive method of imputation. Using Epiclomal's predicted clusters, we infer a missing value with the median (or 0 if this is 0.5) of the observed CpG states in the same cluster, for the same locus. If there are no observed CpG states, then we use the median (or 0) of all cells. If there are still no observed states, we use 0.

##### 2.5 Computational complexity

The Epiclomal methods are computationally more expensive than the non-probabilistic methods, but can provide some advantages in better prediction of the number of clusters (see, for example, Figure 5**g** and panel **c** of Supplementary Figures 2 to 8), cell-to-cluster assignments (see V-measure in, for example, Figure 4) or epiclone prevalence prediction (see Figures 3**c** and 5**e**). EpiclomalRegion is more computationally intensive than EpiclomalBasic, but because it imposes more structure into the data modelling it leads to overall better results (see, for example, Supplementary Figures 5**a**, 5**c** and panel **i** of Supplementary Figures 2 to 5).

##### 2.6 Implementation

Epiclomal was implemented in Python 3. The remaining methods (data pre-processing, synthetic data generator, non-probabilistic methods and evaluation measures) were implemented in R 3.5. The computational framework consists of several pipelines run through the snakemake workflow. Our software is available at https://github.com/shahcompbio/Epiclomal.

#### 3 Synthetic data generator

We generate synthetic data assuming the true cluster-specific methylation profiles differ only at certain regions following a phylogenetic tree structure.

For a given set of parameters as described in Table 2 of the main text, we generate a simulated data set of single-cell DNA methylation by doing as follows.

1. We start by generating the $K$ vectors of true hidden methylation states according to the following steps.

   i. For a total number of loci $M$, we generate $R$ regions with balanced sizes sampled from a multinomial distribution of size $M$ and equal probabilities $1/R$;

   ii. Each vector $\boldsymbol{\mu}_{1r}$ containing the probability of a given locus to be methylated in region $r$ of cluster $k = 1$ is generated from a Dirichlet distribution.

   iii. Each entry of the vector of hidden states (methylated or unmethylated) for cluster $k = 1$, $\mathbf{G}_1$, is generated by sampling from a Bernoulli distribution with probability of success and failure given by the $\boldsymbol{\mu}_{1r}$'s.

iv. We now generate the second vector of true methylation states, $\mathbf{G}_2$, by first setting $\mathbf{G}_2 = \mathbf{G}_1$ and then flipping the methylation states of a proportion of loci from a randomly picked region $r$.

v. We generate $\mathbf{G}_k$ for $k = 3, \dots, K$ by first randomly picking an ancestor vector of methylation states $\mathbf{G}_{k-1}$. We set $\mathbf{G}_k = \mathbf{G}_{k-1}$ and then flip the methylation states of a proportion of loci from a randomly picked region $r$, obtaining $\mathbf{G}_k$.

vi. If $\mathbf{G}_k$ is by chance equal to any previously generated vector we discard $\mathbf{G}_k$ and repeat Step v again.

2. We now generate the true vector of cell clustering assignments $\mathbf{Z}$ by sampling each entry from a multinomial distribution of size one and given cluster prevalence probabilities.

3. We finally generate the observed data for each cell $n$ as follows.

   i. If $Z_n = k$, the vector of observed methylation states for cell $n$ is obtained by sampling the methylation state of each locus from a Bernoulli distribution with probabilities of success and failure depending on the true methylation state at that locus, $G_{krl}$, as in Equation (1).

   ii. To account for the presence of missing data we only keep a certain proportion of observations by choosing at random a locus and then keeping a random number (normally distributed with mean of 10 and standard deviation of 2) of observations to right and to the left of this site. We repeat this step many times till we obtain the desired proportion of observed data, which is one minus the desired missing proportion. By doing this procedure we are simulating sequencing reads that when aligned to the genome they cover multiple consecutive CpGs.

### 4 Evaluation measures

#### 4.1 V-measure

V-measure is our main measure of clustering performance given a ground-truth clustering. V-measure assigns a score between 0 (random clustering) and 1 (perfect clustering) to a predicted set of cell-to-cluster assignments, when compared to the ground-truth clustering.

V-measure is calculated as the harmonic mean between homogeneity ($h$) and completeness ($c$): $V = \frac{2hc}{h+c}$. A predicted clustering has homogeneity 1 if all of the predicted clusters contain only data points which are members of a single true class. A predicted clustering result has completeness 1 if it assigns all of those datapoints that are members of a single true class to a single predicted cluster. Exact formulae of these measures are given in Rosenberg and Hirschberg [12]. We used the Python package scikit-learn to calculate V-measure.

##### 4.2 Number of clusters

The number of predicted clusters is sometimes reported as an additional evaluation measure and compared with the ground-truth or expected number of clusters. This measure does not evaluate how correctly the cells are grouped together. Therefore, it is possible for a prediction to result in the correct number of clusters, but with V-measure close to 0. Nevertheless, the number of predicted clusters is a relevant measure to report in applications involving cancer dynamics as it is of great importance to know the approximate number of tumour clones.

##### 4.3 Clone prevalence mean absolute error

To calculate cluster prevalence MAE (mean absolute error), we assign to each cell $n$ its corresponding true and predicted cluster prevalence values $p_n$ and $\hat{p}_n$, respectively, and compute $1/N \sum_{n=1}^{N} |p_n - \hat{p}_n|$, where $N$ is the total number of cells.

The true cluster prevalence value for a given cell $n$ can be given (in the case of the synthetic data simulations) or calculated as $p_n = \sum_{i=1}^{N} I(c_i = c_n)/N$, where $I$ is the indicator variable that returns true (or 1) if the true cluster of cell $i$ $c_i$ is the same as the true cluster of cell $n$ $c_n$. In other words, we calculate the proportion of cells that have the same true clustering assignment as cell $n$ (including $n$).

The predicted cluster prevalence value $\hat{p}_n$ for a given cell $n$ is calculated by averaging over $N$ the estimated clustering assignment parameters $\pi^*_{nk}$, see Section 2.1 of the Supplementary Material, $\hat{p}_n = \sum_{n=1}^{N} \pi^*_{nk}/N$. This is very similar to calculating the proportion of cells within the same predicted cluster, but since the predicted clustering assignments are probabilities, the result will be slightly different.

##### 4.4 Hamming distance

For each cell in a synthetic dataset, we calculate the hamming distance as the proportion of discordant entries between true and estimated vectors of methylation states, which correspond to the true and predicted clusters for that cell. This is done for all CpG loci, including observed and inferred loci. We then report the mean hamming distance over all cells.

##### 4.5 Uncertainty true positive rate

To compute the uncertainty true positive rate (TPR) for Epiclomal predictions, we proceed as follows. First, we look at the Epiclomal's predicted vector of epigenotypes (matrix **G** in Figure 1 of the main text). For each epigenotype, we go through all the regions and build a list of regions that differ between at least two epiclones. Then, we build a list of "uncertain cells", that is, a list of cells that could belong to more than one cluster because of one or more of the different regions is completely missing. For each of the "uncertain cells" we also build a set of "candidate clusters", that is, all the clusters that cell could belong to. All the cells in the "uncertain cells" list should have a posterior clustering assignment probability of less than 1, roughly 1 divided by the number of possible clusters this cell could belong to (if a cell could belong to one of 2 clusters the probability should be roughly 0.5; if it could belong to one of 3 clusters the probability should be roughly 0.33). If this is the case, then the uncertainty with respect to this cell was predicted well and it is considered a true positive. The uncertainty TPR is calculated as the number of true positives divided by the number of uncertain cells. For example, if all the uncertain cells were predicted with the approximate correct posterior probability, the uncertainty TPR is 1. If all the uncertain cells were given a probability of 1, than the uncertainty TPR is 0. Finally, if the number of uncertain cells is 0, then the uncertainty TPR is undefined. This is an ad-hoc way to estimate the uncertainty TPRs and may not be easily extended to more complicated or noisy cases such as for real data, but it gives us a way to evaluate uncertainty for the simpler and more controlled scenarios considered in our simulations.

### 5 Supplementary Tables

Table 1: *Breast cancer xenograft sc-WGBS data from three patients (SA501, SA535 and SA609).* The InHouse data presented in main text Figures 5 and 6 corresponds to cells from all plates except plate px0837 (xenograft passage 2), summing 558 cells in total. The data presented in the main text Figure 7 corresponds to all SA501 plates, totalizing 244 cells.

| Patient tumour ID | Tumour type | Plate ID | Passage | Number of cells |
|---|---|---|---|---|
| SA501 | TNBC | px0582 | X10A | 45 |
| SA501 | TNBC | px0680 | X10A | 61 |
| SA501 | TNBC | px0443 | X7A | 48 |
| SA501 | TNBC | px0472 | X7A | 50 |
| SA501 | TNBC | px0837 | X2 | 40 |
| SA532 | ER+PR-Her2+ | px0544 | X6 | 68 |
| SA532 | ER+PR-Her2+ | px0650 | X6 | 71 |
| SA609 | TNBC | px0738 | X6 | 48 |
| SA609 | TNBC | px0739 | X6 | 52 |
| SA609 | TNBC | px0740 | X6 | 60 |
| SA609 | TNBC | px0741 | X6 | 55 |

Table 2: An excel spreadsheet (SupplementaryTable2.xlsx) containing the raw coordinates of the 94 CpG Islands considered in Figure 7 of the main text.

Table 3: An excel spreadsheet (SupplementaryTable3.xlsx) containing the annotation of 94 regions considered in Figure 7 of the main text.

#### References

[1] Sébastien A Smallwood, Heather J Lee, Christof Angermueller, Felix Krueger, Heba Saadeh, Julian Peat, Simon R Andrews, Oliver Stegle, Wolf Reik, and Gavin Kelsey. Single-cell genome-wide bisulfite sequencing for assessing epigenetic heterogeneity. *Nature methods*, 11(8):817–820, 2014.

[2] Christof Angermueller, Stephen J Clark, Heather J Lee, Iain C Macaulay, Mabel J Teng, Tim Xiaoming Hu, Felix Krueger, Sébastien A Smallwood, Chris P Ponting, Thierry Voet, et al. Parallel single-cell sequencing links transcriptional and epigenetic heterogeneity. *Nature methods*, 13(3):229, 2016.

[3] Tadeusz Caliński and Jerzy Harabasz. A dendrite method for cluster analysis. *Communications in Statistics-theory and Methods*, 3(1):1–27, 1974.

[4] Jiarui Ding, Sohrab Shah, and Anne Condon. densitycut: an efficient and versatile topological approach for automatic clustering of biological data. *Bioinformatics*, 32(17):2567–2576, 2016.

[5] Ryan M Mulqueen, Dmitry Pokholok, Steven J Norberg, Kristof A Torkenczy, Andrew J Fields, Duanchen Sun, John R Sinnamon, Jay Shendure, Cole Trapnell, Brian J O'Roak, et al. Highly scalable generation of dna methylation profiles in single cells. *Nature Biotechnology*, 2018.

[6] Tony Hui, Qi Cao, Joanna Wegrzyn-Woltosz, Kieran O'Neill, Colin A. Hammond, David J.H.F. Knapp, Emma Laks, Michelle Moksa, Samuel Aparicio, Connie J. Eaves, Aly Karsan, and Martin Hirst. High-resolution single-cell dna methylation measurements reveal epigenetically distinct hematopoietic stem cell subpopulations. *Stem Cell Reports*, 2018.

[7] Yu Hou, Huahu Guo, Chen Cao, Xianlong Li, Boqiang Hu, Ping Zhu, Xinglong Wu, Lu Wen, Fuchou Tang, Yanyi Huang, et al. Single-cell triple omics sequencing reveals genetic, epigenetic, and transcriptomic heterogeneity in hepatocellular carcinomas. *Cell research*, 26(3):304, 2016.

[8] Andrew Roth, Andrew McPherson, Emma Laks, Justina Biele, Damian Yap, Adrian Wan, Maia A Smith, Cydney B Nielsen, Jessica N McAlpine, Samuel Aparicio, et al. Clonal genotype and population structure inference from single-cell tumor sequencing. *Nature methods*, 13(7):573–579, 2016.

[9] Michael I Jordan, Zoubin Ghahramani, Tommi S Jaakkola, and Lawrence K Saul. An introduction to variational methods for graphical models. *Machine learning*, 37(2):183–233, 1999.

[10] David M Blei, Alp Kucukelbir, and Jon D McAuliffe. Variational inference: A review for statisticians. *Journal of the American Statistical Association*, 112(518):859–877, 2017.

[11] David J Spiegelhalter, Nicola G Best, Bradley P Carlin, and Angelika Van Der Linde. Bayesian measures of model complexity and fit. *Journal of the Royal Statistical Society: Series B (Statistical Methodology)*, 64(4):583–639, 2002.

[12] Andrew Rosenberg and Julia Hirschberg. V-measure: A conditional entropy-based external cluster evaluation measure. In *Proceedings of the 2007 joint conference on empirical methods in natural language processing and computational natural language learning (EMNLP-CoNLL)*, 2007.
