## Supplementary Figures for "Epiclomal: probabilistic clustering of sparse single-cell DNA methylation data"

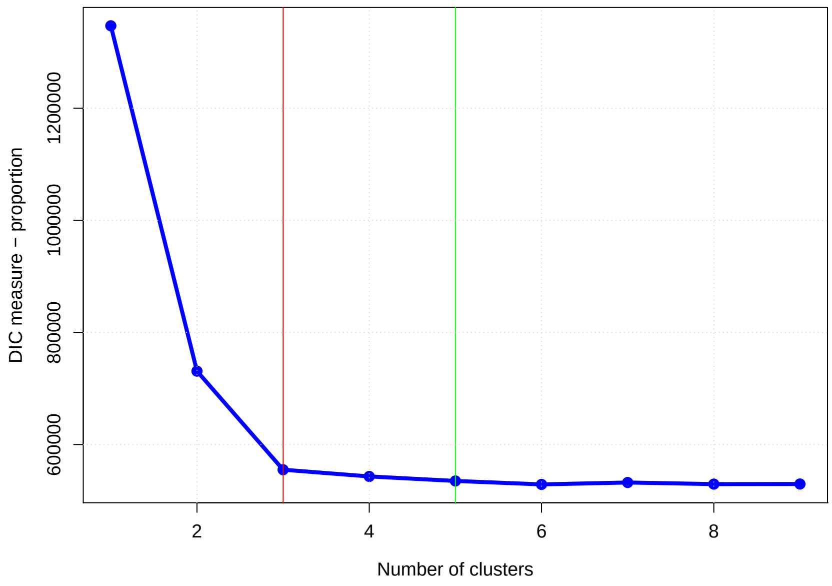

**Supplementary Figure 1** Example of a DIC elbow plot (course-grained) for EpiclomalRegion for the InHouse data set with 10 000 loci. Our DIC algorithm selects only number of clusters for which there is a decrease of at least 2% in DIC values, green vertical line. Then, the elbow value is picked, red vertical line.

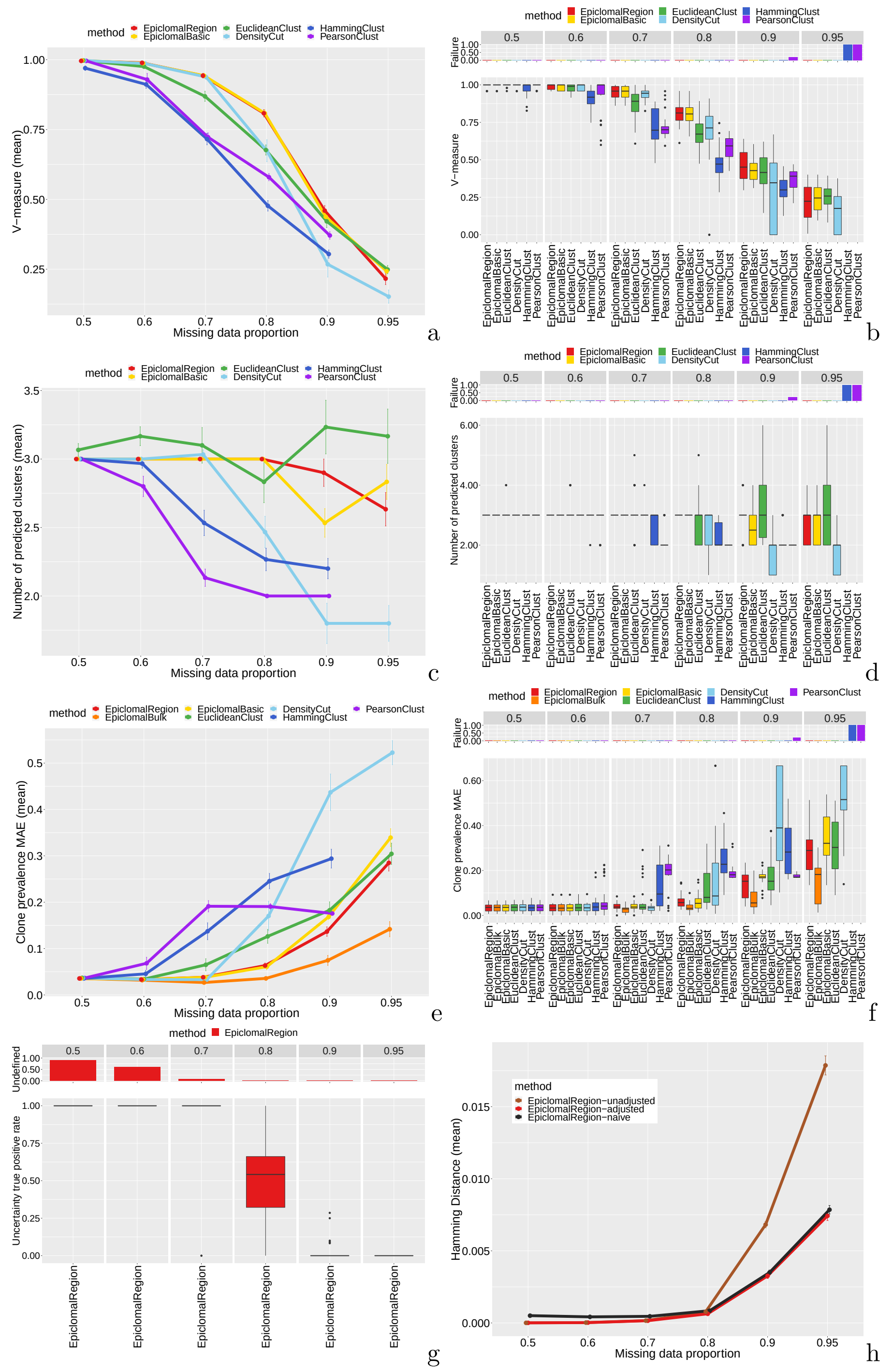

**Supplementary Figure 2 Simulation results when varying the missing proportion.** Panels a and b show results for V-measure in two different ways: means and error bars in panel a, and boxplots in panel b. The barplots above the boxplots show the proportion of data sets for which a method failed to produce a result. Equivalently, panels c and d show results for the number of predicted epiclones. The correct number of clusters is 3. Panels e and f show results for the epiclone prevalence MAE. Panel g shows boxplots for the uncertainty true positive rate (TPR), with the top panel showing the proportion of data sets for which the uncertainty TPR is undefined because there is no true uncertainty, see also Supplementary Material Section 4.5. Panel h shows the average hamming distance for three variants of EpiclomalRegion inferred methylation states: unadjusted, adjusted and naive, see Supplementary Material Sections 2.4 and 4.4.

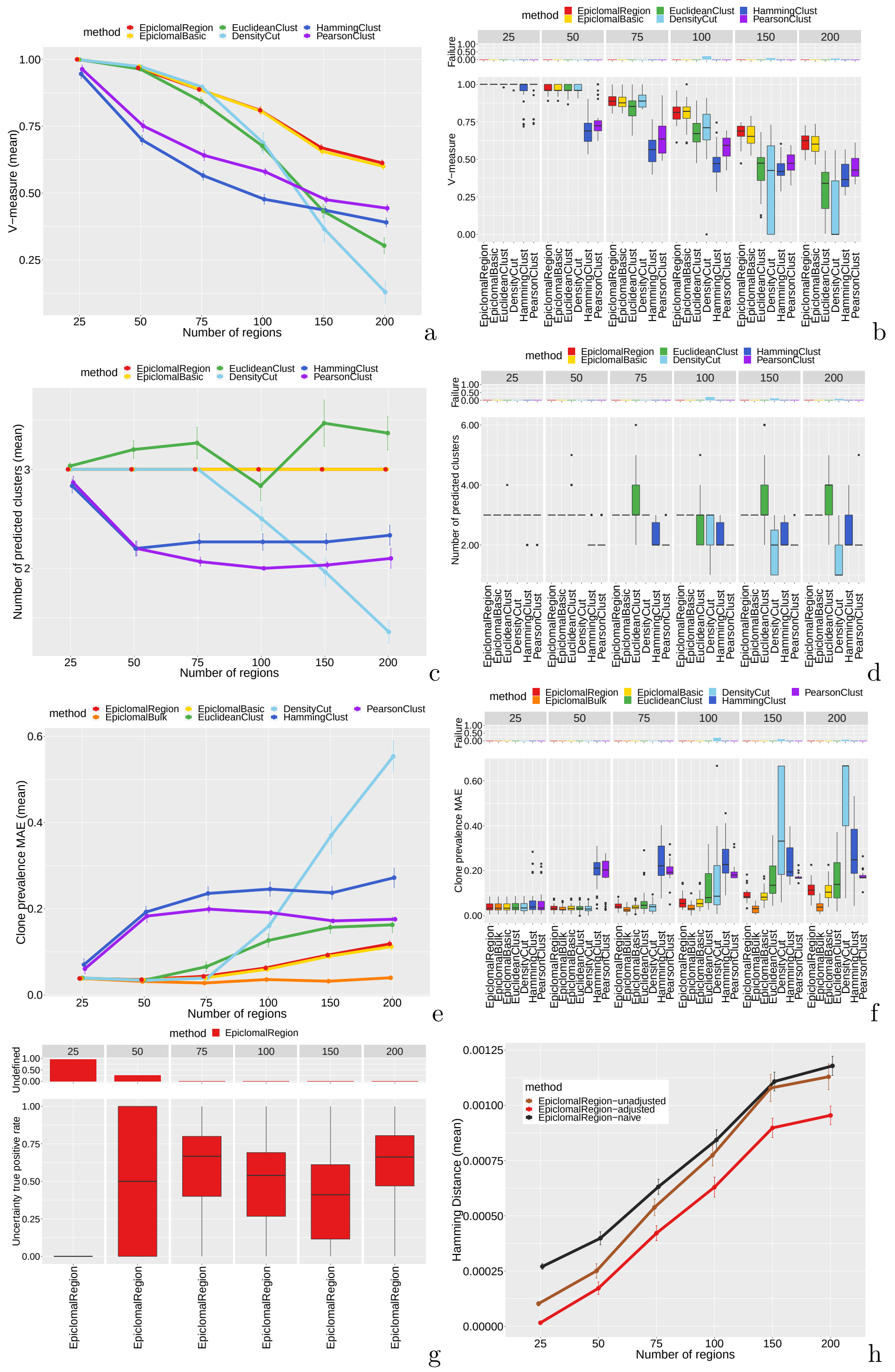

**Supplementary Figure 3 Simulation results when varying the number of regions.** Plots as in Sup. Fig. 2. The larger the number of regions the smaller the differences among epiclones as the number of loci is fixed and our synthetic data generator only allows for one region to change at each new cluster generation. The Epiclomal methods perform better than the other methods and correctly predict the number of clusters (panel c). As expected, all methods perform worse for the largest number of regions because there is less difference between epiclones (200 regions correspond to 0.5% of the loci being different, while 25 regions correspond to 4% of the loci being different). The correct number of clusters is 3.

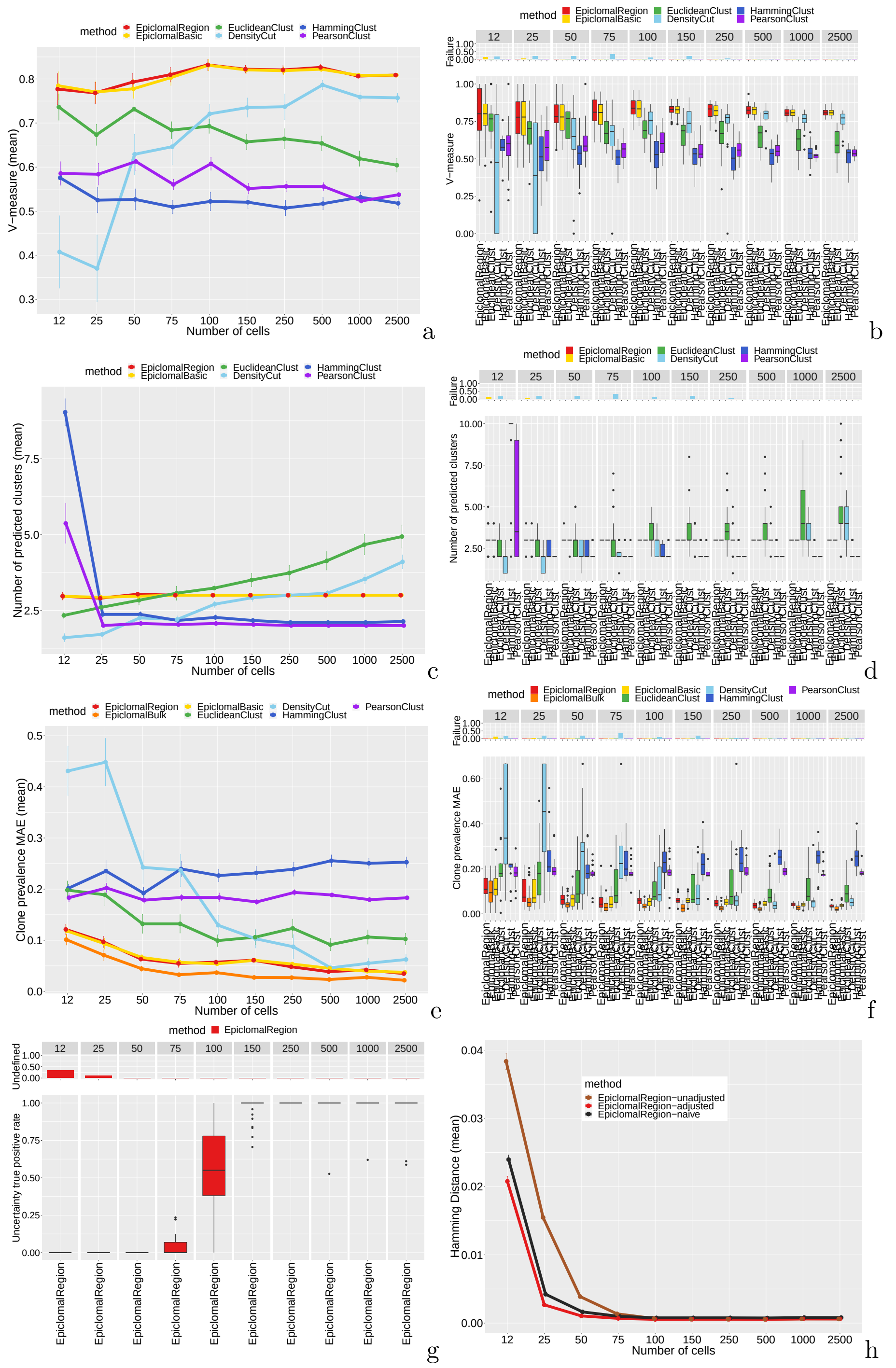

**Supplementary Figure 4 Simulation results when varying the number of cells.** Plots as in Sup. Fig. 2. Increasing the number of cells does not improve the overall V-measure, except for DensityCut, but it reduces the variability of V-measure values. Epiclomal methods produce better V-measures in this case than the other methods. Panel e) shows that EpiclomalBulk produced the best estimates of epiclone prevalences. Starting at about 150 cells EpiclomalRegion was able to obtain an uncertainty true positive rate close to one (panel g). The correct number of clusters is 3.

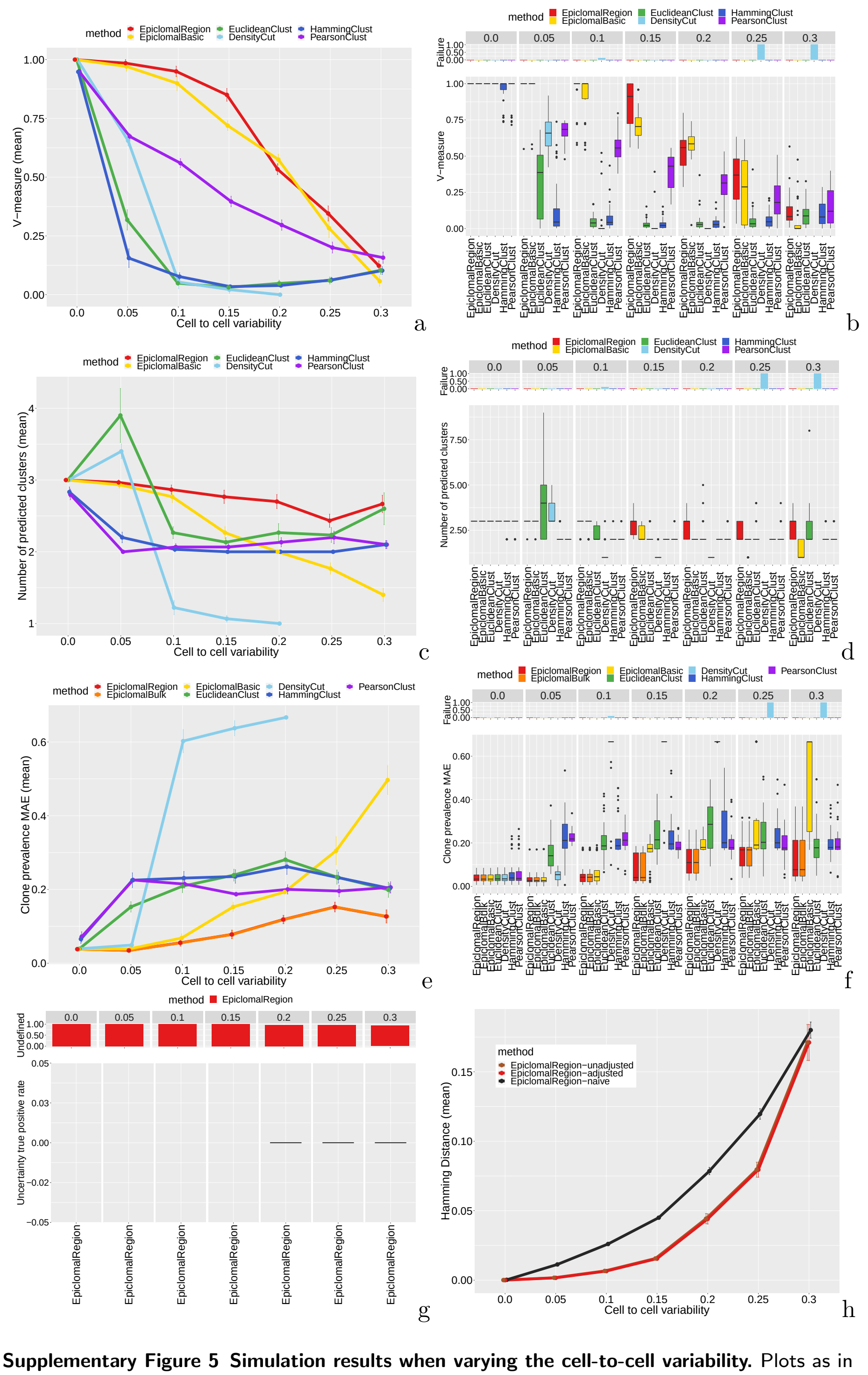

**Supplementary Figure 5 Simulation results when varying the cell-to-cell variability.** Plots as in Sup. Fig. 2. The Epiclomal methods perform significantly better than the other methods when the cell-to-cell variability is between 0.05 to 0.20. A cell-to-cell variability of 0.20 means that at each CpG location that is not in the separating regions (regions that are different among clusters), 20% random cells are in the opposite methylation state than the remaining 80%. Hence a variability of 0.50 means that all the non-separating CpGs have completely random methylation states. When the variability is large ( $\geq 0.25$), all methods perform poorly. The correct number of clusters is 3.

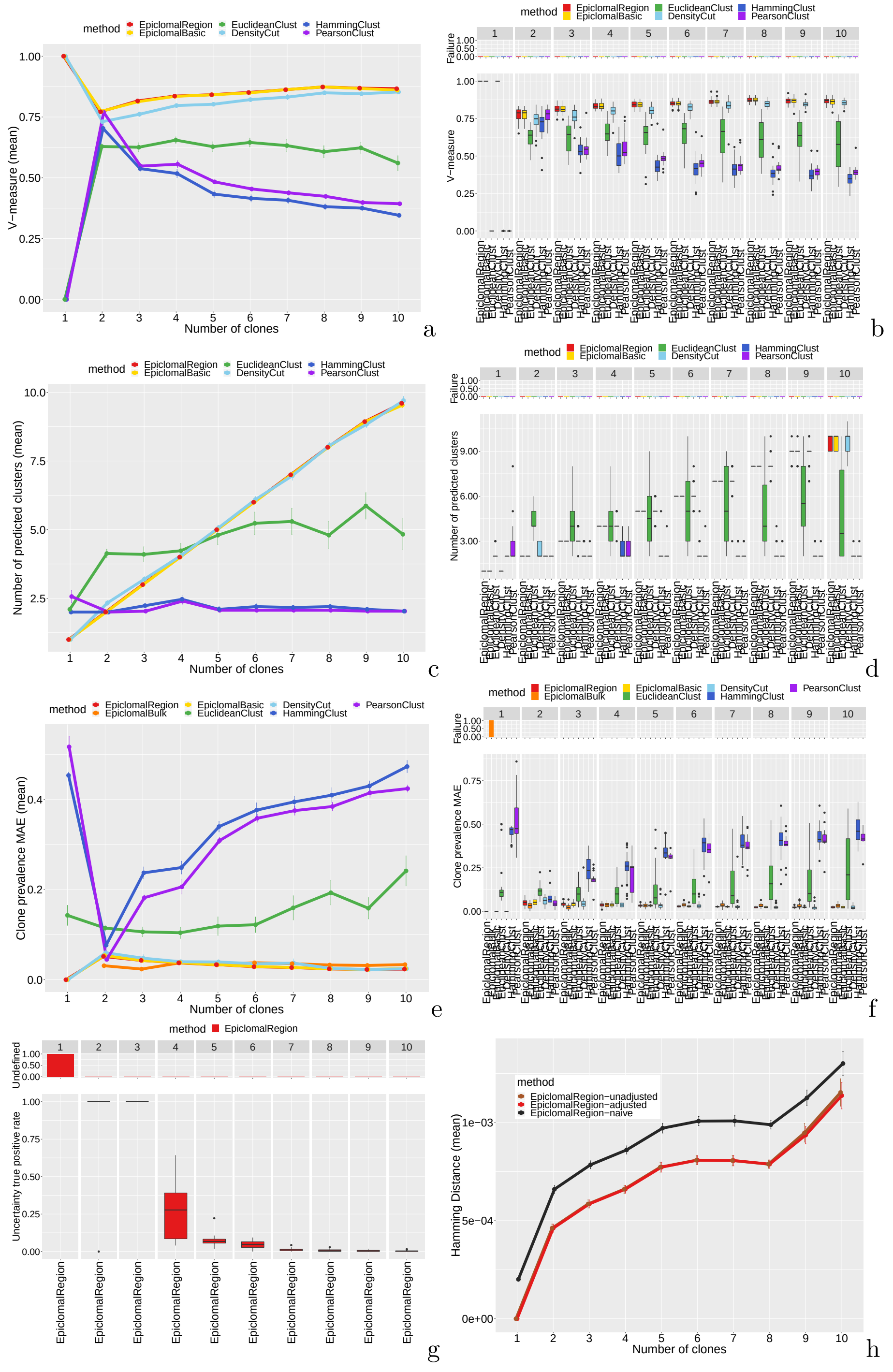

**Supplementary Figure 6 Simulation results when varying the number of epiclones from 1 to 10.** Plots as in Sup. Fig. 2. Epiclomal methods perform slightly better than DensityCut and much better than the other methods in terms of V-measure. Epiclomal and DensityCut are the only methods that can correctly predict the number of clusters (panel c).

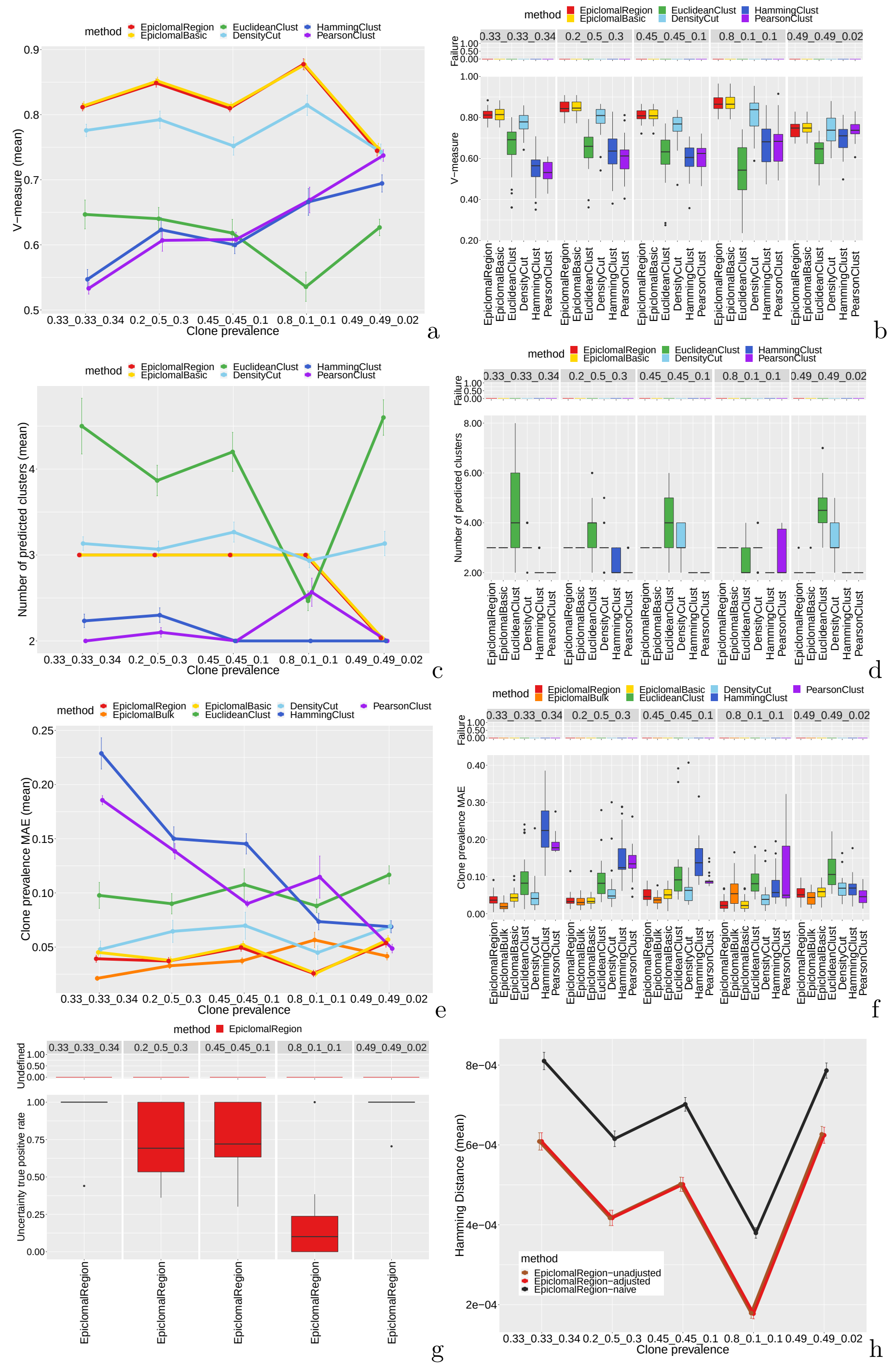

**Supplementary Figure 7 Simulation results when varying the cluster prevalences from equal to very imbalanced.** The number of clusters is three. Plots as in Sup. Fig. 2. EpiclomalRegion and EpiclomalBasic give better V-measures than the other methods, and they correctly predict three clusters, except in the case where one of the clusters has only 2% of the cells. Interestingly, DensityCut does predict three clusters for this difficult case. The uncertainty true positive rate for EpiclomalRegion is above 0.75 for all cases except the case where two of the clusters have only 10% of the cells each.

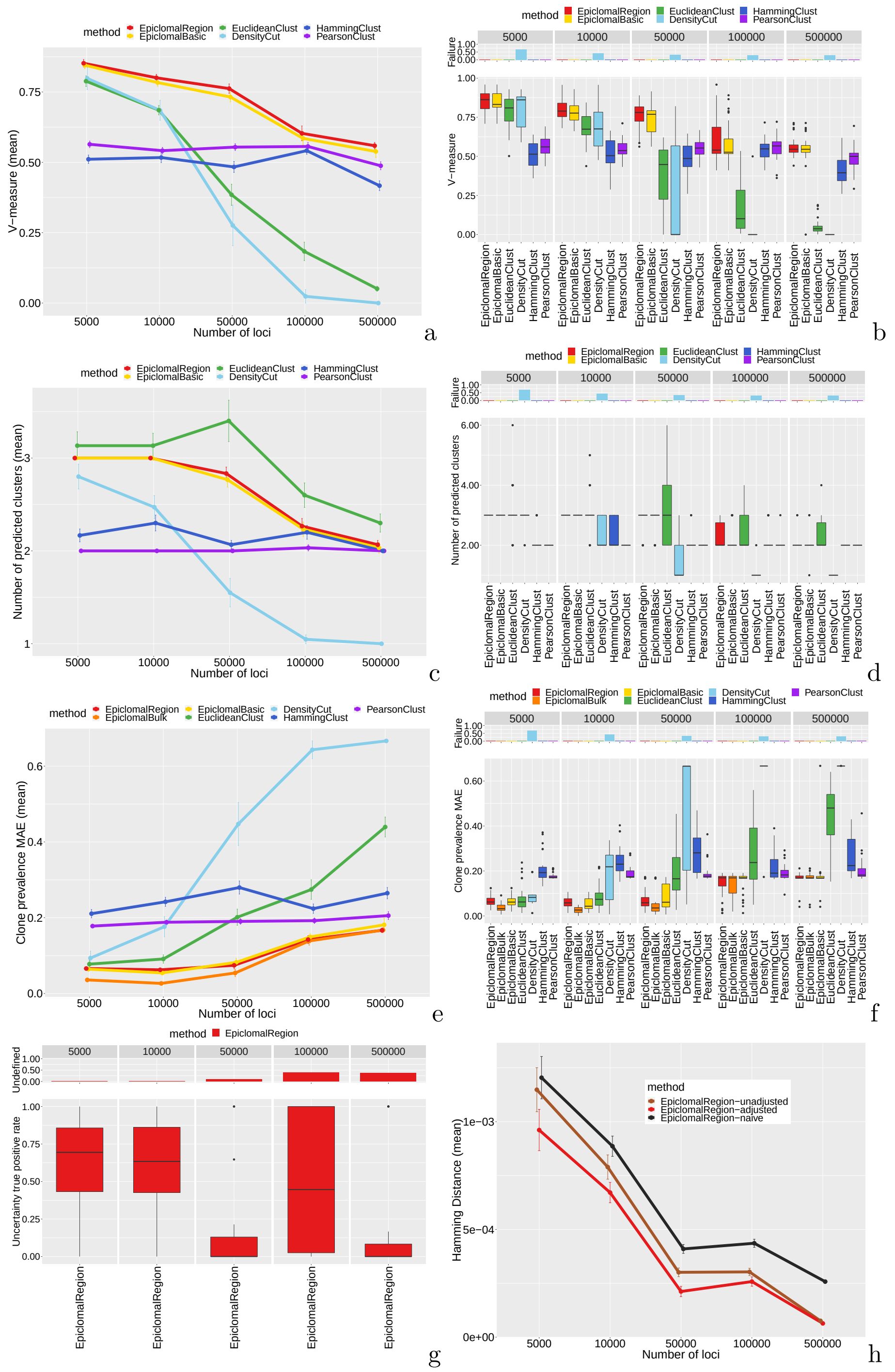

**Supplementary Figure 8 Simulation results when varying the number of loci.** Plots as in Sup. Fig. 2. As we increase the number of loci the number of regions also increase, but their sizes remain fixed and so the amount of CpGs that make the clusters different. The performance of HammingClust and PearsonClust remains somewhat constant as we increase the number of loci, while the other methods show a decreasing pattern in performance. However, the Epiclomal methods still perform better in all cases than all the other methods, especially for 5 000, 10 000 and 50 000 loci. Therefore, this provides support to the strategy of selecting a smaller number of loci (under 50 000) in order to keep the true signal and eliminate noise when analyzing a real data set. The correct number of clusters is 3.

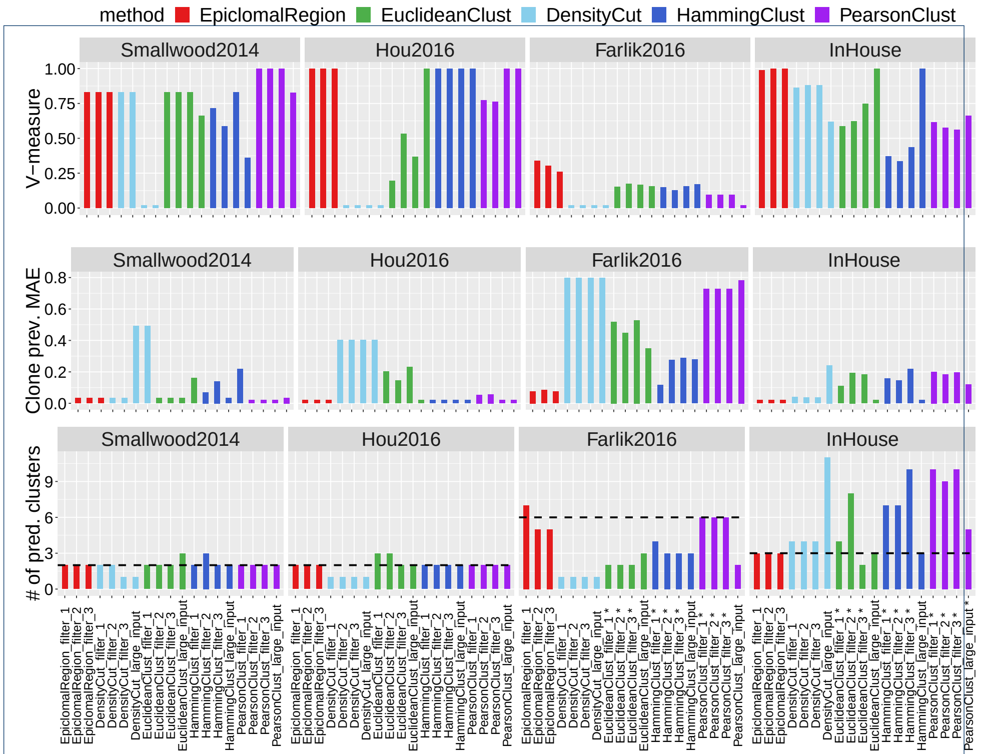

**Supplementary Figure 9** Results for real data sets. (a) V-measure, (b) epiclone prevalence mean absolute error (MAE) and (c) number of predicted clusters for three filtered scenarios for each data set: filter 1, 2 and 3 corresponding to 10 000, 15 000 and 20 000 loci, respectively. For all data sets, we also present the results for a large input data set, in which only the regions with average IQR $< 0.01$ were eliminated. * The stars following the x-axis labels correspond to the cases where a method failed to return a result, in which case we first pre-imputed the missing values with mean region-based values or median CpG-based values across cells and reran the same method, see also Supplementary Material Section 1.5. The true number of clusters are marked by horizontal dashed lines in panel c. See also Table 2. In (b) we consider as true epiclone prevalences the cluster proportions calculated from the clustering reported in the corresponding papers.

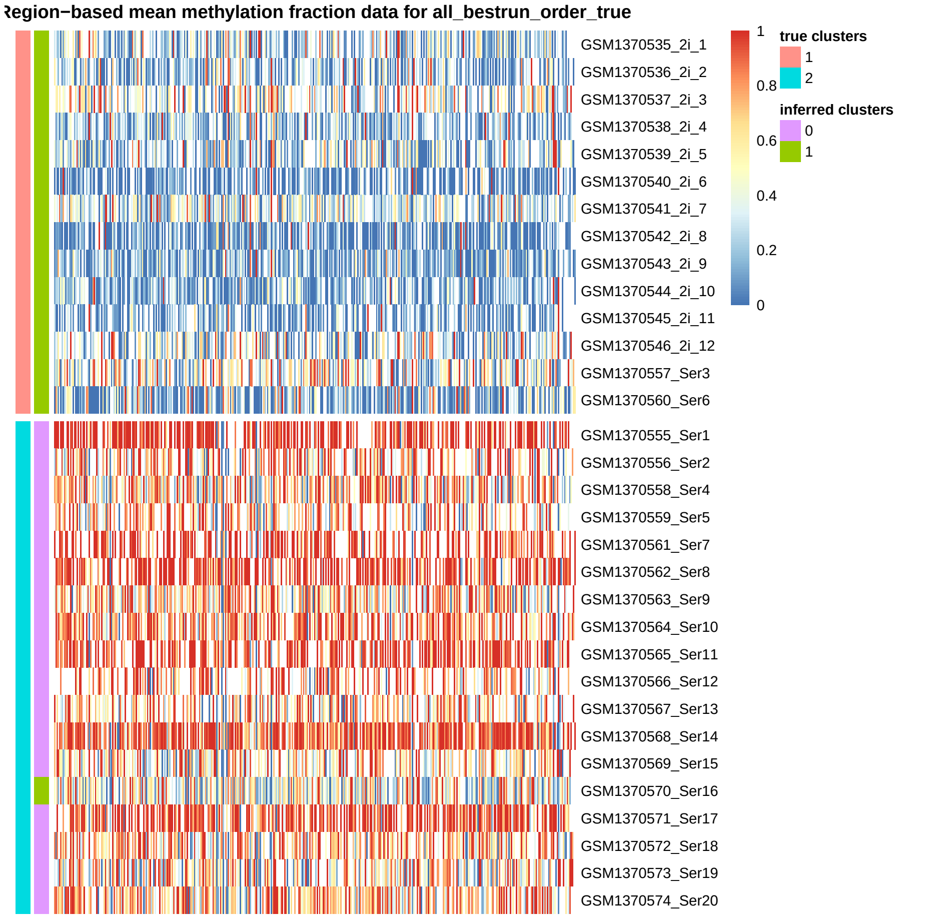

**Supplementary Figure 10** EpiclomalRegion clustering on the Smallwood2014 data set, filtered to include the most variable regions and obtain $\approx$ 10 000 loci (421 regions, average missing proportion 0.69, 32 cells). EpiclomalRegion returned the correct number of clusters (2), but missclassified one cell (GSM1370570_Ser16), V-measure = 0.83. The configuration with the misclassified cell obtained a slightly better DIC score than the ground truth.

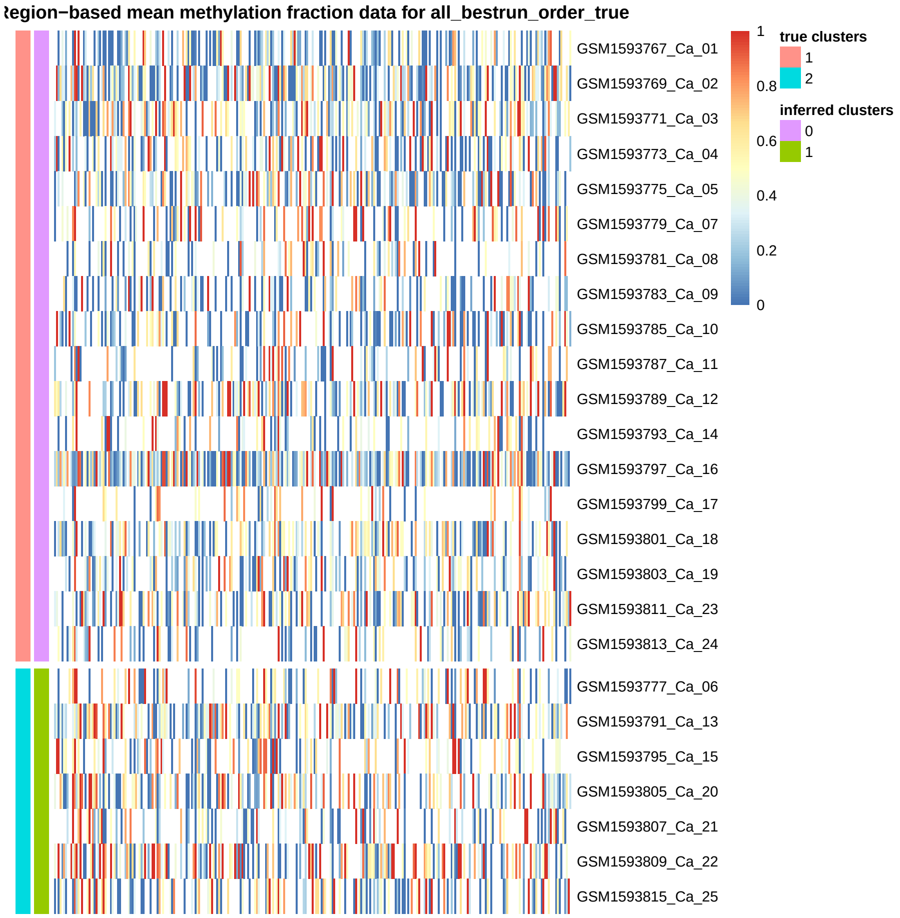

**Supplementary Figure 11** EpiclomalRegion clustering on the Hou2016 data set, filtered to include the most variable regions and obtain $\approx 10\,000$ loci (287 regions, average missing proportion 0.87, 25 cells). EpiclomalRegion correctly classified all cells, V-measure = 1.

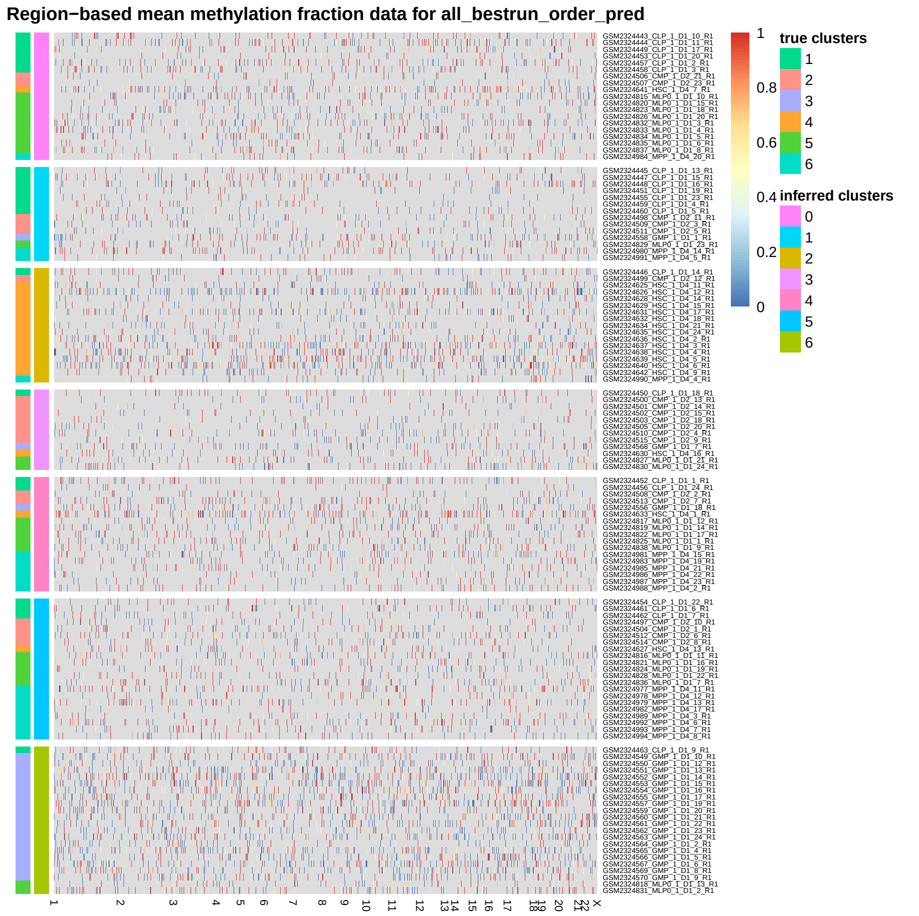

**Supplementary Figure 12** EpiclomalRegion clustering on the Farlik2016 data set, filtered to include the most variable regions and obtain ≈ 10 000 loci. For this set, we eliminated the regions that had on average more than 98% missing CpGs. EpiclomalRegion obtained 7 clusters (ground-truth is 6) and V-measure 0.36.

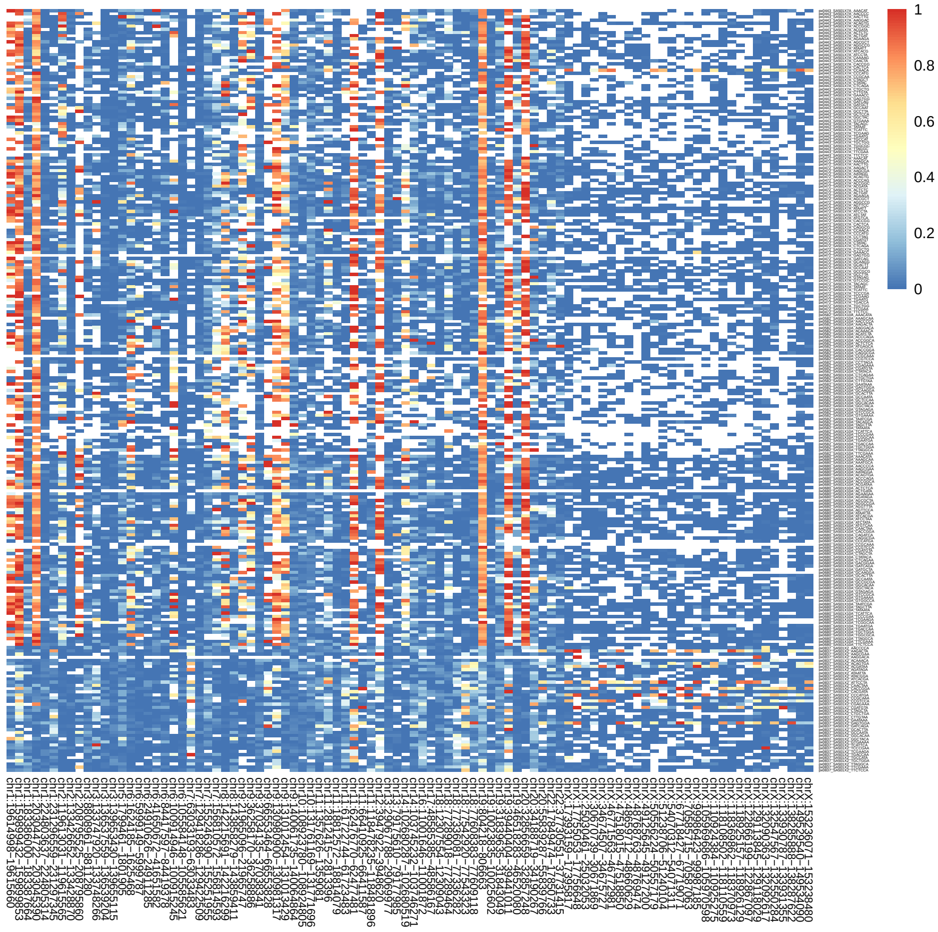

**Supplementary Figure 13** Average methylation for the all the 244 SA501 cells and the 94 NMF-selected regions (CGIs), unclustered. The x-axis shows the region chromosome, start and end.

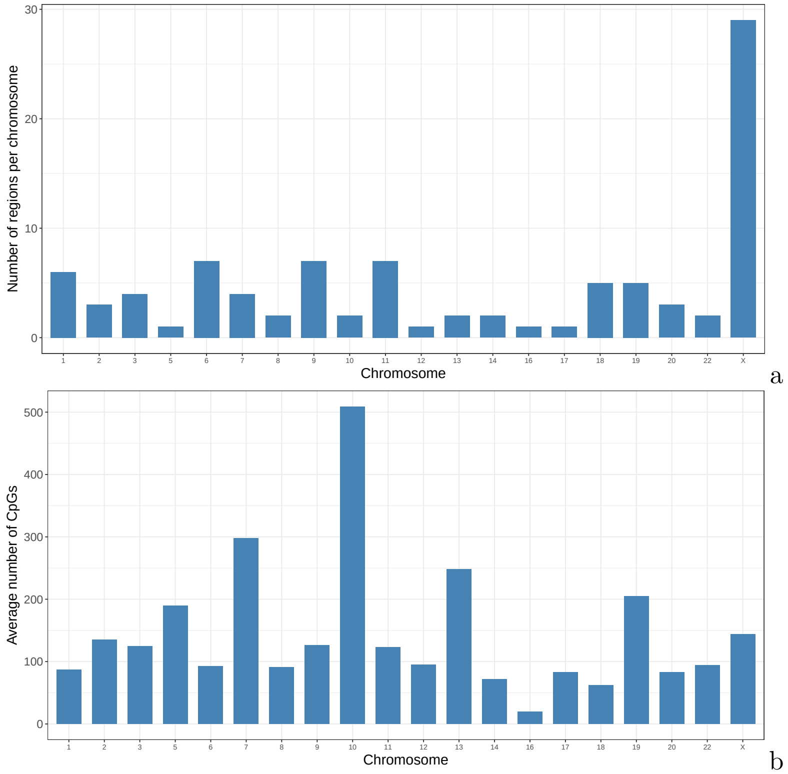

**Supplementary Figure 14** a) Distribution of the 94 NMF-selected regions (CGIs) for SA501 data across the genome. b) Average number of CpGs across regions per chromosome.
